# Autotrophic adaptive laboratory evolution of the acetogen *Clostridium autoethanogenum* delivers the gas-fermenting strain LAbrini with superior growth, products, and robustness

**DOI:** 10.1101/2023.01.28.526018

**Authors:** Henri Ingelman, James K. Heffernan, Audrey Harris, Steven D. Brown, Kurshedaktar Majibullah Shaikh, Asfand Yar Saqib, Marina J. Pinheiro, Lorena Azevedo de Lima, Karen Rodriguez Martinez, Ricardo A. Gonzalez-Garcia, Grant Hawkins, Jim Daleiden, Loan Tran, Hunter Zeleznik, Rasmus O. Jensen, Vinicio Reynoso, Heidi Schindel, Jürgen Jänes, Séan D. Simpson, Michael Köpke, Esteban Marcellin, Kaspar Valgepea

**Author notes:** Correspondence: Esteban Marcellin,; Kaspar Valgepea. These authors contributed equally to this work (Henri Ingelman, James K. Heffernan, Audrey Harris). Present address: Libbs Farmacêutica, 01140-050 São Paulo, Brazil (Marina J. Pinheiro). Present address: Braskem S/A, 13086-530 Campinas, Brazil (Lorena Azevedo de Lima).

## Abstract

Microbes able to convert gaseous one-carbon (C1) waste feedstocks are increasingly important to transition to the sustainable production of renewable chemicals and fuels. Acetogens are interesting biocatalysts since gas fermentation using *Clostridium autoethanogenum* has already been commercialised. However, most acetogen strains need complex nutrients, display slow growth, and are not robust for routine bioreactor fermentations. In this work, we used three different and independent adaptive laboratory evolution (ALE) strategies to evolve the wild-type *C. autoethanogenum* to grow faster, without yeast extract and to be robust in operating continuous bioreactor cultures. Multiple evolved strains with improved phenotypes were isolated on a minimal medium with one strain, named “LAbrini” (LT1), exhibiting superior performance regarding the maximum specific growth rate, product profile, and robustness in continuous cultures. Whole-genome sequencing of the evolved strains identified 25 mutations. Of particular interest are two genes that acquired seven different mutations across the three ALE strategies, potentially as a result of convergent evolution. Reverse genetic engineering of sporulation-related mutations in genes CLAU_3129 (*spo0A*) and CLAU_1957 recovered all three superior features of our ALE strains through triggering significant proteomic rearrangements. This work provides a robust *C. autoethanogenum* strain to accelerate phenotyping and genetic engineering and to better understand acetogen metabolism, which we named “LAbrini”.

## 1. INTRODUCTION

One-carbon (C1) gas (CH_4_, CO_2_, CO) emissions from burning fossil fuels are the main contributors to climate change (1). Alarmingly, this continuing trend might lead to irreversible climate change and a significant loss of biodiversity (2). To tackle and possibly reverse these horrific trends, a circular bioeconomy with decarbonised energy sectors, sustainable chemical and fuel production, and integrated waste management must replace our linear take-make-waste economy (3). While solar and wind are leading the way for decarbonising energy generation, the improvement of various biocatalysts allowing the “up-cycling” of waste C1 feedstocks is considered one of the key strategies for sustainable production of chemicals and fuels towards achieving carbon neutrality (4).

One attractive route toward recycling waste carbon into products is gas fermentation (5). The ideal biocatalysts for this bioprocess are bacteria termed acetogens. Acetogens use the energy-efficient Wood–Ljungdahl pathway (WLP) to fix carbon oxides (CO_2_ and CO) into acetyl-CoA from a wide range of feedstocks (6). Notably, the model-acetogen *Clostridium autoethanogenum* is already used in commercial-scale ethanol production (7) and has been successfully piloted for acetone and isopropanol production (8). However, the lack of non-commercial acetogen strains capable of growing on a minimal medium has slowed the understanding of acetogen metabolism and rational metabolic engineering. The strains currently publicly available uptake gas slowly, grow slowly, require yeast extract (YE), and are challenging to run in continuous bioreactor cultures. Minimal growth medium (e.g. without YE) is required for accurate quantification of carbon distribution and metabolic modelling to understand metabolism and predict phenotypes (9, 10). Faster gas uptake and faster growth would speed up the phenotyping and genetic engineering of acetogens (6, 11). A more robust strain in operating continuous bioreactor cultures (12, 13) would ease generating high-quality steady-state data. Also, the use of various *C. autoethanogenum* strains for academic research complicates cross-comparison between laboratories that using a standard and user-friendly strain could homogenise. A publicly available acetogen strain with the above-mentioned improved features would benefit the academic community enormously.

Adaptive laboratory evolution (ALE) is a powerful and elegant approach for obtaining strains with improved traits. Evolution’s reliance on natural selection to enrich mutants with increased fitness allows strain optimisation without requiring *a priori* knowledge of the genetic alterations necessary to effect such changes (14). Genes that accumulate mutations across multiple ALEs are under stronger selection pressure and serve as attractive targets for further genetic engineering studies (15). Such convergent evolution is a fascinating phenomenon where distinct lineages evolve similar traits or functions independently, often in response to analogous environmental challenges. Observing and comparing these adaptations can shed light on the underlying principles of convergent evolution at the molecular and metabolic levels. ALE has recently been used also for engineering and improving autotrophic growth capabilities. For example, ALE was used to convert the natively heterotrophic workhorse microbes *Escherichia coli* (16) and *Pichia pastoris* (17) to CO_2_-fixing organisms. In acetogens, ALE has improved tolerance towards methanol in *Sporomusa ovata* (18), cyanide in *Clostridium ljungdahlii* (19), benzene, toluene, and xylenes in *C. autoethanogenum* (20), and acetate in *Clostridium* sp. AWRP (21). ALE was also recently used in *Clostridium carboxidivorans* to expand its product spectrum and improve growth on CO_2_+H_2_ (22) and in *Eubacterium limosum* for higher CO tolerance and faster growth (23, 24). Additionally, *C. ljungdahlii* and *C. autoethanogenum* have been adapted by continuous bioreactor cultivation to grow without YE (25, 26), but it is not clear if the phenotype was the result of physiological adaptation (i.e. reversible) or evolution (i.e. permanent).

In this work, we used three different and independent ALE strategies across three continents to obtain superior *C. autoethanogenum* strains that can grow faster, without YE on minimal medium, and are robust for operating continuous bioreactor cultures. As a result, multiple evolved strains with improved phenotypes were isolated, with one strain – named “LAbrini” (LT1) – exhibiting superior performance in terms of maximum specific growth rate (μ_max_), product profile, and robustness in bioreactor cultures. Genome sequencing revealed 25 mutations with two genomic “hot-spot” regions across the evolved strains, and reverse genetic engineering of sporulation-related mutations recovered all three superior features of our ALE strains. This work provides the robust *C. autoethanogenum* strain LAbrini to the academic community to accelerate phenotyping and genetic engineering and facilitate the generation of high-quality steady-state datasets. Additionally, our work showcases the synergistic benefits of ALE, mutation analysis, and reverse engineering for shaping superior gas-fermenting strains.

## 2. MATERIALS AND METHODS

### 2.1. Starting bacterial strain for adaptive laboratory evolution (ALE)

The starting strain for all three ALEs (see next section) was *Clostridium autoethanogenum* strain JA1- 1 (27) deposited in the German Collection of Microorganisms and Cell Cultures (DSMZ) as DSM 10061 and stored as a glycerol stock at ‒80°C.

### 2.2. Three adaptive laboratory evolution (ALE) workflows

#### 2.2.1. University of Tartu (UT) ALE workflow – bottle ALE on CO

Glycerol stock of DSM 10061 was revived in PETC-MES medium (28) supplemented with 5 g/L of fructose, 1.5 g/L of YE, and 0.4 g/L of cysteine-HCl·H_2_O as the reducing agent. Cells were sub-cultured from exponential growth phase into the same medium but with fructose replaced by CO as the carbon source: this culture was Round 1 of UT ALE (Fig. 1a). Cells were then sub-cultured from exponential growth phase to the same medium with CO (Round 2). Starting from Round 3, each round was carried out in 2‒4 replicate bottles. The culture with the fastest growth was sub-cultured to the next round from exponential growth phase. During Rounds 3‒5, YE was removed from the medium stepwise: first to 0.5 g/L (Round 3), then to 0.05 g/L (Round 4), and no YE starting from Round 5 of UT ALE. Serial sub-culturing was stopped after 35 rounds when the decrease of maximum specific growth rate (μ_max_) during UT ALE had levelled off (Fig. 1b).

**Fig. 1.**
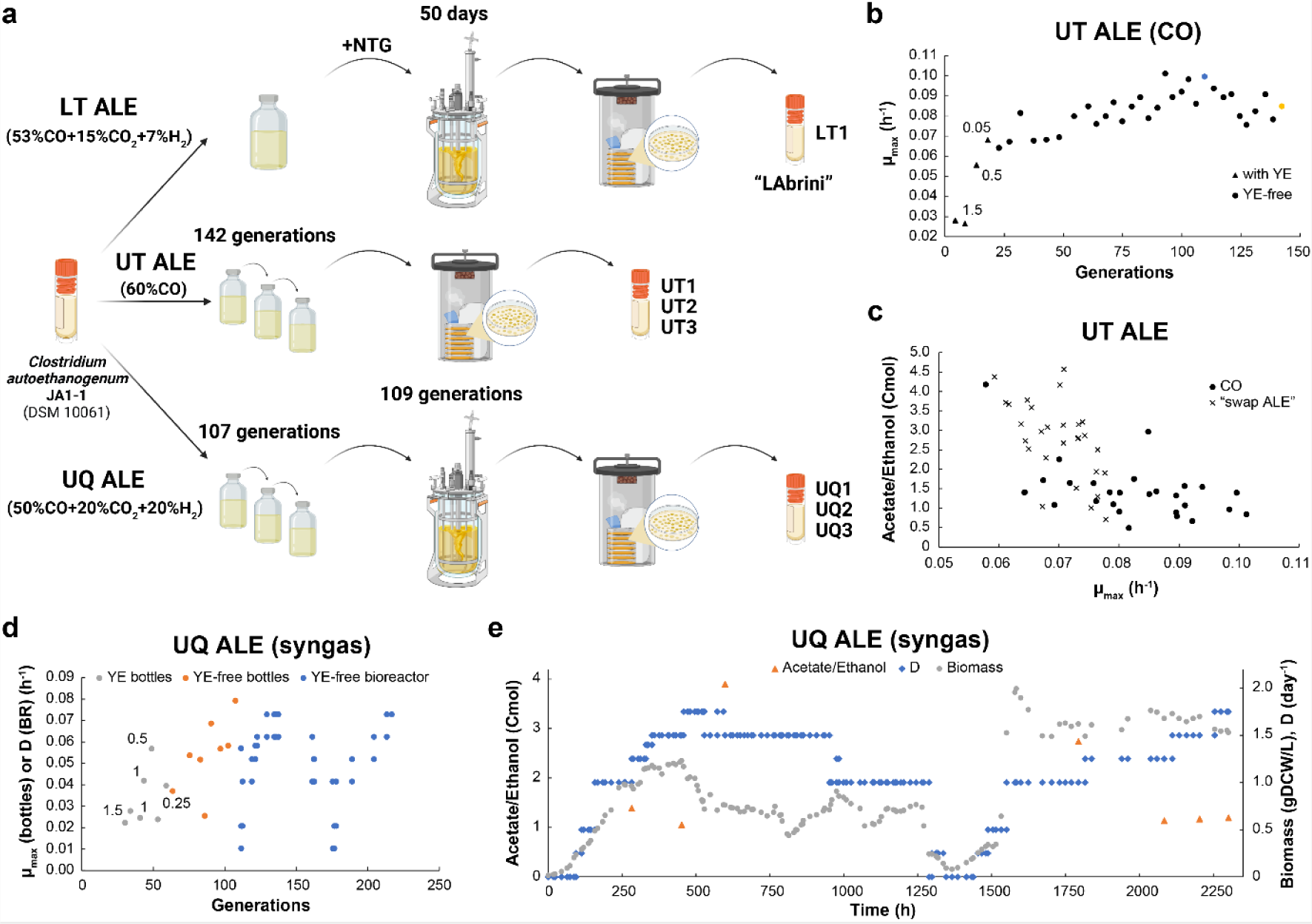
Autotrophic adaptive laboratory evolution (ALE) of *Clostridium autoethanogenum* JA1-1 for improved growth phenotypes executed with three strategies across three facilities. (a) Overview of the different ALE strategies employed. See text for details. Created with BioRender.com. (b) Change of maximum specific growth rate (μmax) with batch culture generations during UT ALE on CO. Blue and yellow circles denote cultures used for isolating single colonies (see text for details). For cultures with generations ∼43, ∼48, ∼96, and ∼105, μmax was calculated using two data points from the exponential phase instead of using 3‒5 data points as for the rest. (c) Change of acetate/ethanol (Cmol) with μmax in combined batch data from UT (UT ALE and UT “swap ALE”; see text for details). (d) Change of μmax in batch or dilution rate (D) in bioreactor (BR) continuous cultures with culture generations during UQ ALE on syngas. (e) Change of acetate/ethanol, biomass concentration (biomass), and D over fermentation time during the continuous culture of UQ ALE on syngas. Numbers next to triangles (panel b) or circles (panel d) denote yeast extract (YE) concentrations. gDCW, gram of dry cell weight.

Bottle cultivations were carried out with 50 mL medium in 250 mL Schott bottles incubated horizontally at 37°C with orbital shaking at 120 RPM under strictly anaerobic conditions. Culture bottle headspace was pressurised to 140 kPa with a 60% CO and 40% Ar gas mixture (AS Eesti AGA). Additionally, a “swap ALE” was performed where the culture was transferred between the previous CO mixture and syngas (50% CO, 20% CO_2_, 20% H_2_, 10% Ar; AS Eesti AGA) (Supplementary Fig. 1). Growth was tracked by measuring culture optical density (OD) at 600 nm. Maximum specific growth rate (μ_max_) was calculated using 3‒5 data points from the exponential phase (except for a few cultures using two data points indicated in Fig. 1b) that yielded a correlation coefficient R^2^ > 0.98 between culture time and natural logarithm (ln) of OD. At the time of sub-culturing, cultures were sampled for extracellular metabolome analysis and occasionally cryo-preserved in glycerol.

Glycerol stocks made from Round 26 (110 generations; blue marker in Fig. 1b) and 35 (142 generations; yellow marker in Fig. 1b) were used to obtain single colony isolates by plating the mixed populations on Round 1 medium with agar (12 g/L) and incubation at 37°C under autotrophic conditions (same gas mixture as for UT ALE). Ten colonies were screened in bottles under the same conditions as in Round 5 of UT ALE (e.g. CO without YE), and three isolates named UT1 (from Round 26) and UT2 and UT3 (from Round 35) were selected for further characterisation (see below) based on μ_max_, or ethanol or 2,3-butanediol (2,3-BDO) yields per biomass (see Results for details). The latter yields (mmol of product/gram of dry cell weight [gDCW]) were estimated by linear regression between the respective product (mmol/L) and biomass concentrations (gDCW/L) that were measured during exponential growth phase with correlation coefficients R^2^ > 0.95.

#### 2.2.2. University of Queensland (UQ) ALE workflow – bottle and bioreactor ALE on syngas

Glycerol stock of DSM 10061 was revived as in Round 1 of UT ALE, and the bottle was pressurised to 125 kPa with CO_2_. Cells were sub-cultured in single replicate bottles from the exponential growth phase into the same medium but with a gradual reduction of fructose (i.e. 2.5, twice 1, and 0 g/L), with 190 kPa syngas (50% CO, 20% CO_2_, 20% H_2_, 10% Ar; BOC Australia) replacing the carbon source (Series 1 of UQ ALE). Next, YE was gradually removed from the medium over 8 bottles (i.e. twice 1.5, twice 1, 0.5, twice 0.25, and 0 g/L). The first YE-free culture was cryo-preserved before pausing the ALE experiment (Series 2 of UQ ALE; grey markers in Fig. 1d). The latter glycerol stock was revived in the same medium as in Round 1 of UT ALE and bottle pressurised to 140 kPa with syngas before sub-culturing in YE-free medium 3-times with 190 kPa and 4-times with 140 kPa syngas (Series 3 of UQ ALE; orange markers in Fig. 1d).

Bottle cultivations were carried out with either 25 or 100 mL medium (125 or 500 mL bottles) and incubated horizontally at 37°C with orbital shaking at 200 RPM under strictly anaerobic conditions. At the time of sub-culturing, strains were occasionally cryo-preserved. Growth was tracked by measuring culture OD at 600 nm and μ_max_ was calculated using three data points from the exponential phase that yielded a correlation coefficient R^2^ > 0.99 between culture time and ln of OD.

The final culture of Series 3 was used as the inoculum for a bioreactor ALE experiment, where the culture was grown on syngas predominantly under chemostat operation at pH 5 (29). Other parameters (e.g. dilution rate, agitation, gas flow rates) were varied based on culture growth (Series 4 of UQ ALE; blue markers in Fig. 1d). After obtaining steady-states at high biomass concentrations and ∼2,300 h of fermentation, the bioreactor culture was plated on solid medium (Round 1 of UT ALE medium with 12 g/L agar) and incubated at 37°C under autotrophic conditions to obtain single colonies. Initially, pure CO at 70 kPa was used, but since only a few colonies appeared after 12 days, the headspace was replaced with syngas, and plates were incubated for an additional 7 days, after which 12 colonies were screened in bottles under the same conditions as at the end of Series 2 and 3 of UQ ALE (e.g. syngas without YE). Three isolates named UQ1, UQ2, and UQ3 were selected for further characterisation (see below) based on μ_max_, or ethanol or 2,3-BDO yields per biomass (see Results for details). The latter yields (mmol of product/gDCW) were estimated with individual replicate (single colony revival) product (mmol/L) and biomass concentrations (gDCW/L), measured during the mid-/late-exponential growth phase.

#### 2.2.3. LanzaTech (LT ALE) workflow – chemical mutagenesis and bioreactor ALE on syngas

Glycerol stock of DSM 10061 was revived in the same medium as in Round 1 of UT ALE and cultured using anaerobic techniques as described previously (8). Chemical mutagenesis was performed by growing cells anaerobically in bottles at 37°C with shaking until mid-exponential growth phase and then exposing cells to the chemical mutagen N-methyl-N’-nitro-N-nitrosoguanidine (NTG) as described before (30), except that after treatment cells were washed with anaerobic medium lacking YE and then transferred and cultured in a chemostat. Bioreactor chemostat gas fermentation with *C. autoethanogenum* has been described before (8) and for selection and adaptation *via* gas fermentation no YE was added to the medium. The culture was grown for approximately 50 days to a final dilution rate (D) of 1 day^−1^ on the gas mixture of 53% CO, 7% H_2_, 15% CO_2_, and 25% N_2_ before single colony isolation and glycerol stock preparation.

### 2.3. Characterisation of single colonies from evolved cultures

#### 2.3.1. Bottle experiments

We carried out additional bottle experiments to more accurately characterise the single colonies isolated from the evolved cultures. We used the same conditions as for Round 5 in UT ALE (e.g. CO without YE) for UT ALE isolates and LT1 (UT1, UT2, UT3, LT1) and the same conditions as for Series 2 in UQ ALE (i.e., syngas without YE) for UQ ALE isolates and LT1 (UQ1, UQ2, UQ3, LT1). We also cultured the ALE starting strain JA1-1 for comparison with YE (Round 2 conditions in UT ALE) and without YE (Round 5 conditions in UT ALE) on CO and syngas with UT ALE cultivation conditions. These cultures were sampled frequently from the exponential phase to enable a more accurate estimation of μ_max_ and product yields. μ_max_ was calculated using 3‒12 data points from the exponential phase that yielded a correlation coefficient R^2^ > 0.99 between culture time and ln(OD). Product yields for UT isolates, LT1, and JA1-1 (mmol of product/gDCW) were estimated as described in section 2.2.1 with correlation coefficients R^2^ > 0.94. For UQ isolates and LT1 on syngas, product yields were calculated from the difference of two time points in mid-exponential growth.

#### 2.3.2. Bioreactor chemostat cultures

Cells were grown either on CO (60% CO and 40% Ar; AS Eesti AGA) for UT1, UT2, UT3, and RE2 (see below) or syngas (50% CO, 20% H2, 20% CO2, and 10% Ar; AS Eesti AGA) for UQ1 and LT1 in a chemically defined medium without YE as described before (29). Cells were grown under strictly anaerobic conditions at 37°C and at pH 5 maintained by 2.5 M or 5 M NH_4_OH. Chemostat continuous cultures were performed in 1.4 L Multifors bioreactors (Infors AG) at a working volume of 750 mL connected to a Hiden HPR-20-QIC mass spectrometer (Hiden Analytical) for online high-resolution off-gas analysis. The system was equipped with peristaltic pumps; mass flow controllers (MFCs); pH, oxidation-reduction potential (ORP), and temperature sensors. Antifoam (435530; Sigma-Aldrich) was continuously added to the bioreactor at 10 μL/h to avoid foaming.

Chemostat cultures were run at D ∼0.5, ∼1, and ∼2 day^−1^ (μ∼0.02, ∼0.04, and ∼0.08 h^−1^) across various isolates with variable gas-liquid mass transfer (gas flow rate and agitation) to maintain similar steady-state biomass concentrations (Table 1). In total, 33 biologically independent steady-states were achieved after OD, gas uptake and production rates had been stable (<15% variability) for at least three working volumes. Bioreactor off-gas analysis using a mass spectrometer for determination of specific gas uptake (CO and H_2_) and production rates (CO_2_ and ethanol) (mmol/gDCW/day) and determination of carbon recoveries and balances have been described before (29).

**Table 1.** Summary of 33 steady state chemostat cultures of UT, LT1, and UQ1 isolates.

| Strain | Feed gas | Gas flow rate (mL/min) | Agitation (RPM) | D (day<sup>-1</sup>) | Biomass concentration (gDCW/L) | Number of replicates |
| --- | --- | --- | --- | --- | --- | --- |
| UT2 | CO | 35 | 690 | 1.00 ± 0.00 | 0.52 ± 0.03 | 3 |
| UT3* | CO | 28 | 570 | 0.49 ± 0.01 | 1.03 ± 0.15 | 3 |
| UT1 | CO | 28 | 570 | 0.53 ± 0.00 | 1.13 ± 0.06 | 3 |
| UT2 | CO | 24 | 570 | 0.52 ± 0.01 | 0.88 ± 0.09 | 4 |
| LT1 | Syngas | 50 | 675 | 0.99 ± 0.04 | 1.41 ± 0.09 | 4 |
| LT1 | Syngas | 72 | 800 | 1.97 ± 0.11 | 1.50 ± 0.07 | 4 |
| LT1 | Syngas | 35 | 525 | 0.50 ± 0.06 | 0.42 ± 0.05 | 2 |
| UQ1 | Syngas | 35 | 525 | 0.50 ± 0.06 | 0.47 ± 0.05 | 2 |
| RE2 | CO | 21 | 570 | 0.50 ± 0.02 | 0.97 ± 0.12 | 4 |
| RE2 | CO | 45 | 680 | 0.96 ± 0.01 | 1.66 ± 0.07 | 4 |

### 2.4. Biomass concentration analysis

Biomass concentration in gDCW/L was determined by measuring culture OD at 600 nm and using the correlation coefficient between culture OD and DCW established at 0.23 (UT and LT ALE isolates) or 0.21 (UQ ALE isolates) using the methodology described before (31).

### 2.5. Extracellular metabolome analysis

Analysis of exo-metabolome was performed using filtered broth samples stored at −20°C until analysis. Organic acids and alcohols were analysed by HPLC (Shimadzu Prominence-I LC-2030 plus system) using a Rezex™ ROA-Organic Acids H+ (8%) 300 ×7.8 mm column (00H-0138-K0; Phenomenex) and a guard column (03B-0138-K0; Phenomenex). Twenty microlitres of the sample were injected using an auto-sampler and eluted isocratically with 0.5 mM H_2_SO4 at 0.6 mL/min for 30 min at 45°C. Compounds were detected by a refractive index detector (RID-20A; Shimadzu) and identified and quantified using relevant standards using the software LabSolution (Shimadzu). We note that cells produced 2R,3R-butanediol.

### 2.6. Whole-genome sequencing (WGS) and mutation analysis

We performed whole-genome sequencing (WGS) for isolated clones UT1, UT2 and UT3 (UT ALE); UQ1, UQ2, and UQ3 (UQ ALE); LT1 (LT ALE); and also the ALE starting strain JA1-1 (DSM 10061) to ensure the accurate determination of mutations that appeared during ALE experiments. Genomic DNA for sequencing was extracted from exponential phase cultures of UT and UQ isolates using the MasterPure Gram Positive DNA Purification Kit (MGP04100; Biosearch Technologies) or Qiagen DNeasy Powersoil Pro Kit (47016; Qiagen) following the manufacturer’s instructions. Genomic DNA for the LT1 strain was extracted with a phenol-chloroform extraction protocol. Purified DNA was quantified for UT and UQ isolates using the NanoDrop^TM^ 1000 instrument (Thermo Scientific) or the Qubit 2.0 instrument (Invitrogen). DNA sequencing was performed using the MiSeq v2 Micro 300 cycles sequencing kit (MS-103-1002; Illumina) on the MiSeq sequencer (Illumina) with 2 × 151 bp paired-end dual indexed (2 × 8 bp) reads or using the Nextera DNA Flex Library Preparation Kit (20018705; Illumina) on the NovaSeq6000 sequencer (Illumina) with 2 x 150 bp paired-end reads. For LT1, DNA processing and sequencing was outsourced to third parties to generate 150 bp pair-end reads using the NovaSeq 6000 Illumina platform and continuous long reads from the PacBio Sequel II instrument, which were used to produce a genome assembly (see below).

WGS data analysis, including trimming, filtering, mapping assembly, and annotation, was carried out using the tool breseq version 0.37.0 (32) with default settings for detecting missing coverage evidence and new sequence junctions and following parameters: limit fold coverage, 500; require match fraction, 0.95; polymorphism frequency cutoff, 1; polymorphism reject indel homopolymer length, 0; polymorphism reject surrounding homopolymer length, 0; polymorphism minimum variant coverage each strand, 1; polymorphism prediction base quality cutoff, 30; polymorphism minimum total coverage, 2; and polymorphism minimum total coverage each strand, 1. With paired-end read overlaps ignored, the total genome coverage was 50.78 for LT1, 410.67‒551.11 for UQ isolates, 88.35‒92.70 for UT isolates, and 75.30 for JA1-1. Mean mapping quality was 58.99 for LT1, 58.95‒59.07 for UQ isolates, 59.32‒59.34 for UT isolates, and 59.3274 for JA1-1. High-quality reads were mapped against the NCBI reference genome assembly ASM148472v1 (GenBank CP012395.1) for *C. autoethanogenum* JA1-1 (33). Two mutations were excluded from further analysis as they were considered non-relevant: one mutation was present in all isolates and the starting strain JA1-1, and one mutation was present in JA1-1 only.

Additionally, for LT1, the Illumina data was used to polish the PacBio genome assembly with the Pilon v1.24 tool (34). The polished assembly was deemed to be highly accurate based on the observation that no 100% frequency variants were predicted by breseq v0.35.1 (32) using the polished assembly and Illumina data as inputs. The validated LT1 assembly was then deposited to the NCBI genome submission portal (Genbank CP110420) for annotation with the NCBI Prokaryotic Genome Annotation Pipeline. See Supplementary Table 1 for a comparison of strain LT1 (LAbrini) gene identifiers (IDs or locus_tags) with previously published complete assemblies of *C. autoethanogenum* JA1-1 genome sequences (33, 35).

### 2.7. Transcriptome analysis of UT isolate bioreactor chemostat cultures

Transcriptome analysis for the steady-state chemostat cultures of UT isolates listed in Table 1 was performed using RNA sequencing (RNA-seq). In addition, the transcriptome of the non-steady-state culture of UT2 at D = 1 day^−1^ without apparent “CO toxicity” (end of culture in Supplementary Fig. 2; see text below for details) was also determined. This enabled the comparison of the transcriptome of the “CO toxicity” steady-state culture of UT2 at D = 1 day^−1^ with a “non-CO toxicity” culture at the same D.

Ten millilitres of culture were pelleted by centrifugation (5,000 × *g* for 3 min at 4°C) and resuspended in 5 mL of RNAlater (76106; Qiagen). Samples were stored at 4°C overnight, centrifuged (4,000 × *g* for 10min at 4°C), and pellets stored at −80°C until RNA extraction. Thawed pellets were resuspended in RLT buffer (74104; Qiagen) containing β-mercaptoethanol and lysed with acid-washed glass beads (G4649; Merck) using the Precellys^®^ 24 instrument with liquid nitrogen cooling (Bertin Technologies). Total RNA was extracted using the RNeasy mini kit (74104, Qiagen) with off-column TURBO^TM^ DNase treatment (AM2239; Invitrogen), followed by purification and enrichment using the RNA Clean and Concentrator^TM^ kit (R1018, Zymo). The efficiency of the total RNA purification and DNA removal was verified using the NanoDrop^TM^ 1000 instrument (Thermo Scientific) and the quality of RNA extracts was checked using the TapeStation 2200 equipment (Agilent Technologies). Total RNA concentration was determined using the Qubit 2.0 instrument (Q32866; Invitrogen). Next, ribosomal RNA (rRNA) was removed using the QIAseq FastSelect –5S/16S/23S Kit (335925; Qiagen) and stranded mRNA libraries were prepared using the QIAseq Stranded RNA Lib Kit (180743; Qiagen). RNA sequencing was performed using the NextSeq MID150 sequencing kit (20024904; Illumina) on the NextSeq500 sequencer (Ilumina) with 2 x 75 bp paired-end dual indexed (2 x 8 bp) reads.

The quality of raw NextSeq reads was verified using MultiQC (36) and adapter sequences were trimmed using the Cutadapt Python package (version 2.10; (37)) allowing a minimum read length of 35 nt. The resulting high-quality paired-end reads were mapped to the NCBI reference genome assembly ASM148472v1 (GenBank CP012395.1) for *C. autoethanogenum* JA1-1 (33) using the align function within Rsubread package (version 2.4.2; (38)). Then, four .bam files per sample were merged using Samtools (version 1.10; (39)) and genomic features were assigned using the featureCounts functions within Rsubread. Transcript abundances were determined as described before (29). Genes were determined to be differentially expressed by fold-change > 1.5 and q-value < 0.05 after false discovery rate correction (FDR) (40) (Supplementary Table 2). RNA-seq data has been deposited in the NCBI Gene Expression Omnibus repository under accession number GSE225123.

### 2.8. Reverse genetic engineering of mutations in JA1-1

The detailed steps of our CRISPR/Cas9-aided homologous recombination workflow for genetic engineering include designing and selecting sgRNAs, plasmid construction and electroporation into *C. autoethanogenum* JA1-1, screening and plasmid curing, and confirmation of transformants were described before (28). Briefly, the homology arms (HAs) for the CLAU_3129 (*spo0A*) deletion plasmid were obtained by PCR amplifying the 1 kb upstream and downstream regions of the *spo0A* gene. The derived 5′ and 3′ HA fragments were fused using overlap extension PCR and moved into the backbone plasmid to create an intermediate plasmid pGFT104. Two unique sgRNAs targeting the *spo0A* gene were selected and cloned into plasmid pGFT104 using a quick-change PCR protocol (41), resulting in plasmids pGFT106 and pGFT107. Nine transformant colonies were screened altogether and the selected isolate named RE1. A similar approach was employed for constructing the plasmid (pGFT088) for generating the SNP in CLAU_1957 (mutation #10 in UT1), wherein the HAs arms were obtained by PCR amplifying the 1kb upstream and downstream regions of the target nucleotide and the SNP was incorporated into the HAs using the quick-change PCR protocol. Only one sgRNA was identified and used where the SNP was part of the PAM sequence, thus disrupting it after homologous recombination to prevent Cas9 from cleaving the target site again. Seven transformant colonies were screened and the selected isolate was named RE2. Supplementary Table 3 lists the primers used and Supplementary Table 4 the strains and plasmids used or constructed in the genetic engineering part of the current study. See Supplementary File 1 (pGFT106), Supplementary File 2 (pGFT107), and Supplementary File 3 (pGFT088) for plasmid maps.

Growth characterisation of the reverse engineered strains RE1 and RE2 was carried out as described in section 2.3. For autotrophic bottle cultures, strains were grown using same conditions as for Round 5 in UT ALE and for RE1 additionally on syngas. Correlation coefficient R^2^ values for μ_max_ and product yield calculations were in the same limits as stated in section 2.3.1 with the exception of the acetate yield for one RE1 CO culture being R^2^ = 0.9.

### 2.9. Proteome analysis of reverse engineered strains and JA1-1

Proteome analysis was carried out from three bioreplicate cultures for each of the two reverse engineered strains RE1 and RE2, and JA1-1. Strains were grown using UT ALE cultivation conditions on syngas and medium of Round 2 in UT ALE (with YE). Cultures were sampled at OD∼0.4 and μ_max_ was calculated using 4‒5 data points from the exponential phase that yielded a correlation coefficient R^2^ > 0.98 between culture time and ln of OD.

Sampling, sample preparation, LC-MS/MS analysis using data-independent acquisition (DIA), and DIA MS data analysis were performed as described before (28). Briefly, cell pellets were treated with a chaotrope-based lysis buffer and bead beating, lysates alkylated and processed, and proteins digested. Peptides were then analysed on a Ultimate 3500 RSLCnano system (Dionex) and a Q-Exactive HF (Thermo Fisher Scientific) mass spectrometer. Analysis of DIA MS data was performed using the DIA-NN software suite (42) and the protein sequence database of assembly ASM148472v1 for *C. autoethanogenum* JA1-1 (NCBI Genbank CP012395.1) (33). This allowed us to confidently quantitate 31,449 peptides and 2,257 proteins across all samples, and 27,030 peptides and 2,152 proteins on average within each sample after removing shared peptides from analysis.

Protein expression fold-changes (FC) with q-values between RE1 and JA1-1, and RE2 and JA1-1 were determined using the software Perseus (43) with Student’s T-test. Only proteins quantified with at least two peptides (2,215 across all samples) were used to ensure higher quantification accuracy. Proteins were considered to be differentially expressed by a FC > 2 and a q-value < 0.05 after FDR correction (40). Differentially expressed proteins are presented in Supplementary Table 5. Proteomics data have been deposited to the ProteomeXchange Consortium (http://proteomecentral.proteomexchange.org) via the PRIDE partner repository (44) with the data set identifier PXD047330.

### 2.10. Sporulation tests for reverse engineered strains and JA1-1

Autotrophic cultures for sporulation tests were the same as described for proteome analysis while heterotrophic cultures were grown in 10 mL of YTF (28) in 50ml serum bottles and incubated horizontally at 37°C with orbital shaking at 120 RPM under strictly anaerobic conditions. Samples from the stationary phase during ∼40 and ∼20 days of autotrophic and heterotrophic incubation, respectively, were observed on glass slips covered with an agarose pad using phase-contrast microscopy (Zeiss Observer Z1) at ×1000 magnification. Heterotrophic cultures were also plated before and after heat-shock (bottles stored for 10 min in a 80°C water bath) on YTF agar (28) and plates were photographed after colonies appeared in 7-8 days.

## 3. RESULTS

### 3.1. Autotrophic adaptive laboratory evolution (ALE) of *Clostridium autoethanogenum* JA1-1 for improved growth phenotypes

Three different autotrophic ALE strategies were employed across three facilities (Fig. 1a; see Materials and Methods for details) to improve the wild-type *Clostridium autoethanogenum* JA1-1 strain’s growth phenotype. For the ALE experiment at the University of Tartu (UT ALE), serial propagation of batch bottle cultures was employed for 142 generations with the gradual removal of YE at the start. For the ALE experiment done at The University of Queensland (UQ ALE), firstly, serial propagation of batch cultures in serum bottles for 107 generations was used with the gradual removal of YE at the start. After that, the evolved population was transferred into a continuous bioreactor culture, where it was grown for another 109 generations without YE. Moreover, for the third ALE experiment done at LanzaTech, Inc. (LT ALE), cells from a bottle culture were exposed to the chemical mutagen NTG after which the culture was transferred into a continuous bioreactor culture and grown without YE for ∼50 days. The UT ALE experiment was conducted on CO, while the LT and UQ ALE experiments were done on syngas (CO, CO_2_, H_2_).

In UT ALE, serial propagation of CO bottle cultures allowed the YE in the media to be removed stepwise by the fourth subculture (Fig. 1b). Notably, a sudden ∼2-fold increase of μ_max_ was observed during YE removal for the third sub-culture (first bottle with reduced YE, 0.5 g/L). The population reached its peak μ_max_ of 0.10 h^−1^ (∼4.6-fold higher compared to the start) by the ∼93^rd^ generation, after which a declining μ_max_ trend was observed (Supplementary Fig. 3). This phenomenon was previously observed when carrying out ALE under stable conditions (45, 46). We also tested if higher μ_max_ values could be reached when transferring the ALE culture between CO and syngas (“UT swap ALE”), but this did not occur (Supplementary Fig. 1). Intriguingly, combined data from UT (UT ALE and “UT swap ALE”) revealed a declining trend of the acetate/ethanol ratio with faster growth (Fig. 1c).

In UQ ALE, the experiment was started with four fructose batch cultures and then transferred to syngas bottles with stepwise removal of YE (Fig. 1d). The increase of μ_max_ during YE removal was not as abrupt as in UT ALE but the increasing trend was clear and the final batch bottle culture with ∼107 generations reached a μ_max_ of 0.08 h^−1^ (∼3.2-fold higher compared to first syngas bottle). Then, the evolved population was transferred to a chemostat bioreactor system where additional selective pressure was applied by changing the dilution rate (D; i.e. μ under steady-state) for ∼109 generations. We obtained a stable culture at high biomass concentrations and D = 1.8 day^−1^ (μ = 0.08 h^−1^) and the UQ ALE bioreactor culture seemed to excrete less acetate compared to ethanol with faster growth above D = 1 day^−1^ (Fig. 1e).

In LT ALE, *C. autoethanogenum* was first treated with the chemical mutagen NTG during growth with YE after which the culture was used to inoculate a continuous bioreactor culture to a final D of 1 day^−1^ without YE for ∼50 days (Fig. 1a). For all the three ALE strategies, ALEs were terminated once significant changes in evolving population phenotypes were no longer observed (e.g. μ_max_ and maximum biomass in bottle cultures, gas data in continuous cultures).

### 3.2. Isolated ALE clones exhibit enhanced growth characteristics and metabolite production patterns

Next, we plated the evolved mixed populations to isolate single clones for detailed characterisation and for evaluating the level of phenotypic heterogeneity within the mixed populations. For UT ALE, we isolated clones from both the culture with peak μ_max_ of ∼0.10 (blue marker in Fig. 1b) and the last ALE bottle with lower μ_max_ (yellow marker in Fig. 1b). Ten colonies were isolated for both UT and UQ ALE, respectively, for preliminary comparison of growth and by-product patterns in batch cultures with the same gas and media as in ALE. Notably, all isolates could be grown without the presence of YE, thus fulfilling the first aim of our work. For UT ALE, we expected that isolates from the peak μ_max_ population (blue marker) would grow faster than isolates from the last slower-growing population (yellow marker): this was not, however, observed (Supplementary Fig. 4a). Variability in μ_max_ of UQ ALE isolate batch cultures was also seen (Supplementary Fig. 4b). Similarly, phenotypic heterogeneity in terms of by-product patterns within isolates of both UT and UQ ALE was observed (Supplementary Fig. 4c-d). In particular, some isolates of UT ALE showed lower acetate/ethanol ratios, while three isolates from UQ ALE with high ethanol production were detected. As a result, three isolates from both UT (UT1, UT2, UT3) and UQ ALE (UQ1, UQ2, UQ3) were selected for further detailed characterisation based on μ_max_ and intriguing metabolite production patterns in the preliminary screen (indicated with asterisks on Supplementary Fig. 4). From LT ALE, one isolate termed LT1 was selected for further study.

We grew the seven selected isolates in autotrophic batch bottle cultures without YE with frequent sampling to more accurately compare growth and by-product patterns. UT and UQ ALE isolates were grown on the gas they were evolved on, CO and syngas, respectively; the isolate obtained through chemical mutagenesis – LT1 – was grown on both. For comparison, the starting wild-type strain for all ALE strategies – *C. autoethanogenum* JA1-1 – was also cultured on both gases but with YE supplementation as we could not observe growth on CO without YE after 520 h. On CO, isolate UT3 had the highest μ_max_ (0.08 h^−1^) among UT ALE while LT1 showed the fastest growth on CO with a μ_max_ of 0.10 h^−1^ (Fig. 2a). In comparison, the starting ALE strain JA1-1 showed a μ_max_ of 0.02 h^−1^ (with YE) that is ∼2- to ∼4-fold lower than all the evolved isolates tested on CO. During growth on syngas, UQ1, UQ3, and LT1 showed similar growth (μ_max_ ∼0.08 h^−1^) that was ∼1.3- and ∼3-fold higher than for UQ2 and JA1-1 (with YE), respectively (Fig. 2b). All evolved isolates grew faster than the starting strain, thus fulfilling the second aim of our work of obtaining faster-growing strains.

**Fig. 2.**
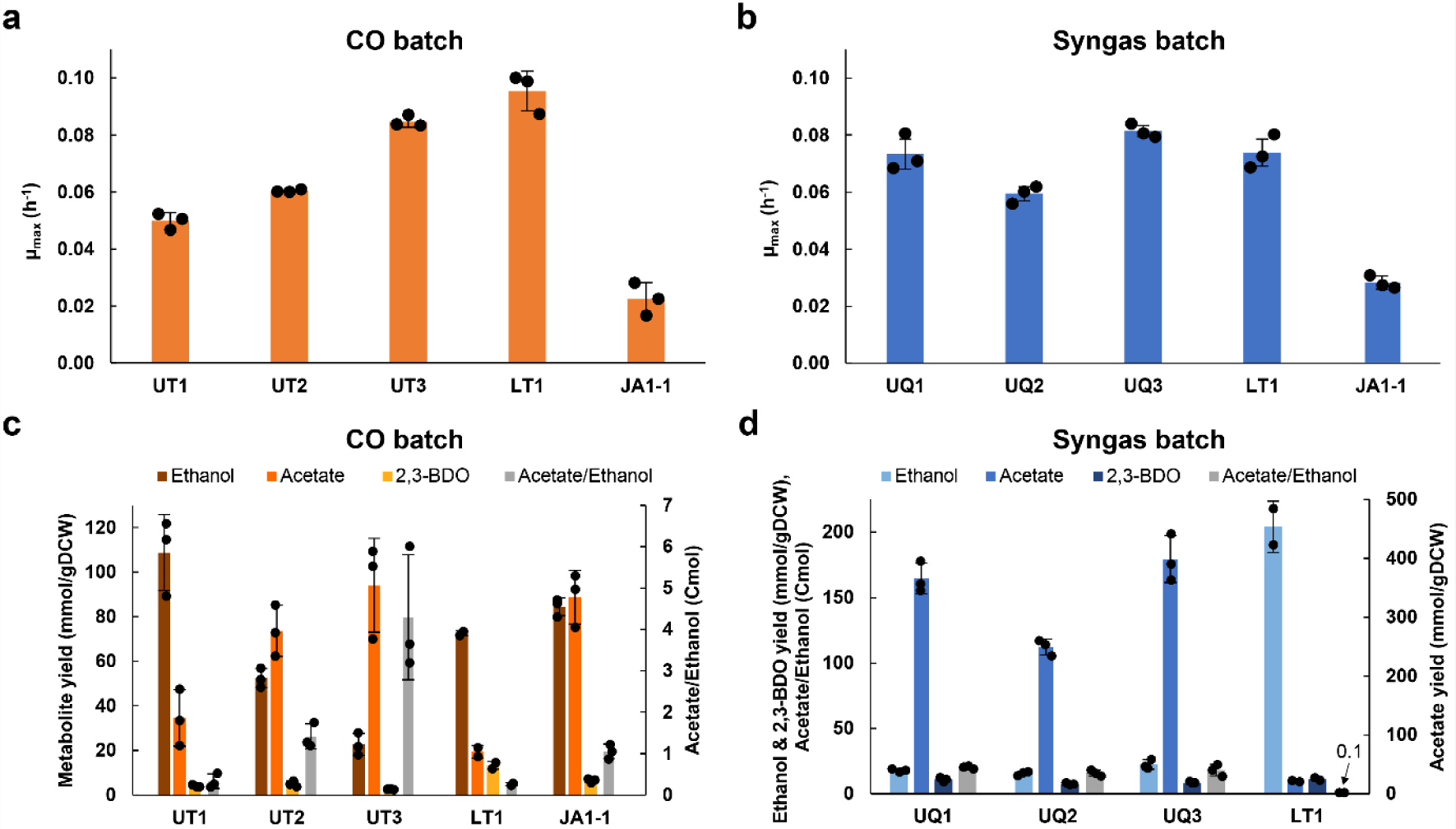
Autotrophic characterisation of selected isolates from UT, UQ, and LT ALE, and the wild-type *C. autoethanogenum* starting strain of ALE (JA1-1) in batch cultures. (a) Maximum specific growth rate (μmax) of UT and LT ALE isolates and JA1-1 in CO batch cultures. (b) μmax of UQ and LT ALE isolate LT1 and JA1-1 in syngas batch cultures. (c) Production yields (mmol/gDCW) of growth by-products and acetate/ethanol (Cmol) of UT and LT ALE isolate LT1 and JA1-1 in CO batch cultures. (d) Production yields (mmol/gDCW) of growth by-products and acetate/ethanol of UQ and LT ALE isolate LT1 in syngas batch cultures. All data for UT, UQ, and LT1 are from yeast extract (YE)-free medium, JA1-1 data are from medium with YE. Bars show average ± standard deviation between bioreplicates, except for LT1 production yields (two bioreplicates). gDCW, gram of dry cell weight; 2,3-BDO, 2,3-butanediol.

Frequent sampling also enabled us to accurately determine production yields (mmol/gDCW) for the detected growth by-products acetate, ethanol, and 2,3-BDO. Notably, the lowest acetate/ethanol ratios were detected on CO for isolates LT1 (0.27) and UT1 (0.32) that were ∼4- and ∼3-fold lower than for JA1-1 (1.05) (Fig. 2c). While UT1 showed the highest ethanol production yield with the lowest μ_max_ of the evolved strains, the fastest growing UT ALE isolate UT3 showed the highest acetate production yield and acetate/ethanol of 4.14 (Fig. 2a and c). The production yield of 2,3-BDO was the highest for LT1, making the strain overall the most favourable on CO in terms of reduced by-products (Fig. 2c). Importantly, LT1 demonstrated preferable metabolite production also during growth on syngas in comparison with UQ isolates (Fig. 2d): it showed a >6-fold higher ethanol production yield with the lowest acetate production yield, translating into a low acetate/ethanol ratio of 0.10. UQ1 and LT1 showed the highest 2,3-BDO production yields. Interestingly, higher acetate production yields for UT and UQ isolates correlate well with higher μ_max_. At the same time, LT1 achieves fast growth with significantly lower acetate production (Supplementary Fig. 5). These extracellular metabolomics data support the notion that evolving acetogens in the presence of C1 substrates trigger rearrangements in central carbon metabolism (18).

### 3.3. Bioreactor continuous gas fermentation cultures show the superiority of isolate LT1

We next tested the performance and robustness of all UT ALE isolates, LT1, and UQ1 in continuous gas fermentation bioreactor as this is the industrially relevant condition (7). Steady-state continuous cultures also provide the necessary data to describe the cell’s physiological state (e.g., μ, gas uptake, by-products, carbon distribution) (47). In total, 33 biologically independent steady-states were achieved across chemostat cultures for strains grown on CO or syngas at D ranging from ∼0.5 to ∼2 day^−1^ (D = µ at steady-state) (Table 1).

We initially aimed to run all strains at D = 1 day^−1^ but failed after multiple attempts to obtain steady-state at D = 1 day^−1^ on CO for UT2 at CO uptake rates (COUR) of ∼1200 mmol/L/day and biomass concentrations of ∼1.5 gDCW/L. We did obtain steady-state at D = 1 day^−1^ for UT2 at COUR ∼400 mmol/L/day, and interestingly CO uptake could only be increased after strongly decreasing the CO% in the bioreactor headspace through changing gas-liquid mass transfer (Supplementary Fig. 2; between ∼380 to ∼440 h), but no steady-state was obtained. Thus UT2 might have struggled due to CO toxicity. For this “CO toxicity” steady-state of UT2 at D = 1 day^−1^ (Table 1; up to ∼380 h on Supplementary Fig. 2), transcriptome analysis revealed a starkly different transcriptional profile compared to all other cultures (Supplementary Fig. 6). The transcriptional profile was also very distinct from the unstable “non-CO toxicity” phase of the same culture operated at same D (from ∼480 h on Supplementary Fig. 2). Notably, we detected strong up-regulation of numerous hypothetical, putative membrane, and sporulation-related proteins and down-regulation of the HytA-E/FdhA complex components (Supplementary Table 2). After these challenges with UT2 cultures and gas uptake with UQ1 at D = 1 day^−1^, we used D = 0.5 day^−1^ to compare strains across gases.

For UT3 though, we could obtain steady-state data at D = 0.5 day^−1^ only regarding gas uptake and production since acetate and ethanol production were highly sensitive to any operational activities to the culture (e.g. halting agitation for a volume check, sampling 1% of culture volume). Furthermore, adjustment of gas-liquid mass transfer was needed even after obtaining target agitation and gas flow (Fig. 3). In striking comparison, operating LT1 cultures was straightforward. Thus we additionally achieved steady-states at D = 1 and 2 day^−1^ at high COUR and biomass concentrations (Fig. 3). The COUR for LT1 growing on syngas at D = 1 day^−1^ is triple of the steady-state value of UT2 on CO (Supplementary Fig. 7). Moreover, faster gas-liquid mass transfer ramping and higher Ds also meant that two steady-states were achieved for LT1 with half the time it took to achieve one steady-state for UT3 (Fig. 3). The robustness of LT1 is highly relevant for the acetogen gas fermentation field as it makes it significantly less challenging to operate continuous gas fermentation cultures to obtain high-quality data. Thus a robust strain for bioreactor continuous gas fermentation was obtained that fulfils the third aim of our work.

**Fig. 3.**
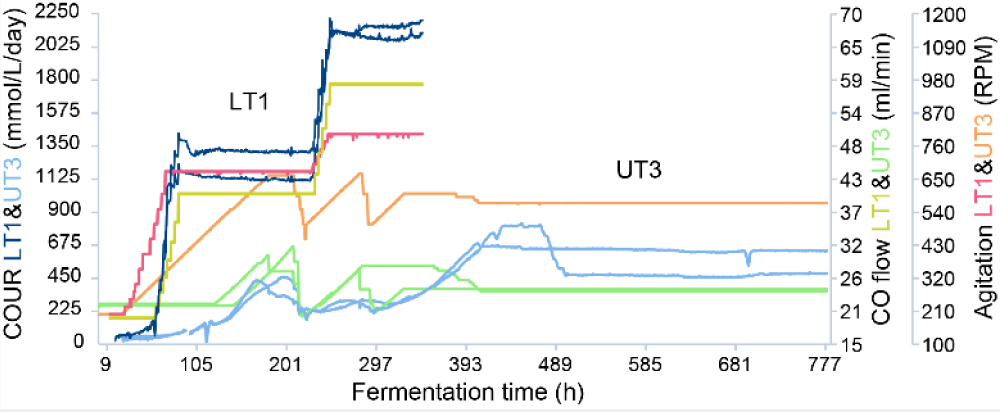
Robustness of strain LT1 in terms of operation of bioreactor continuous gas fermentation cultures. LT1 data are for the chemostats that achieved steady-states at specific growth rates (μ) of 1 and 2 day^−1^. See Table 1 for more details on LT1 and UT3 chemostats. COUR, CO uptake rate (mmol/L/day); RPM, rounds per minute.

A comparison of steady-state data from the bioreactor runs demonstrates that various metabolic rearrangements took place during the three ALE strategies (Fig. 4). Different from the batch data (Fig. 2), UT strains showed similar growth characteristics in CO chemostats (D = 0.5 day^−1^) except for 2,3-BDO production (Fig. 4). While the acetate concentration for UT2 was lower most likely due to lower biomass concentration (Fig. 4a), specific by-product production rates (Fig. 4b) and specific gas rates (Fig. 4c) showed similar values, except for the specific production rate of 2,3-BDO (q_2,3-BDO_; mmol/gDCW/day). Chemostat results were different from batch data also when comparing UQ1 and LT1 in syngas chemostats at D = 0.5 day^−1^ (Fig. 4). The two strains showed very similar performance at the same D both in terms of by-product production and gas uptake and production. Notably, LT1 outperformed all strains concerning acetate/ethanol ratios by showing values <1 at D = 1 and 2 day^−1^ (Fig. 4a). The ability to achieve higher biomass concentrations for LT1 D = 1 and 2 day^−1^ cultures also translated into the highest ethanol and 2,3-BDO levels (Fig. 4a) and due to faster growth also to higher specific production rates (q_EtOH_ and q_2,3-BDO_) (Fig. 4b). LT1 cultures at D = 1 and 2 day^−1^ also show the highest carbon flux to the reduced ethanol and 2,3-BDO products across the various strains and steady-states (Fig. 5). Similarly to the latter strain, the specific H_2_ uptake rate (q_H2_) for LT1 did not increase proportionally with D (Fig. 4c), suggesting another avenue for improving *C. autoethanogenum* performance through ALE, i.e. improving H_2_ uptake.

**Fig. 4.**
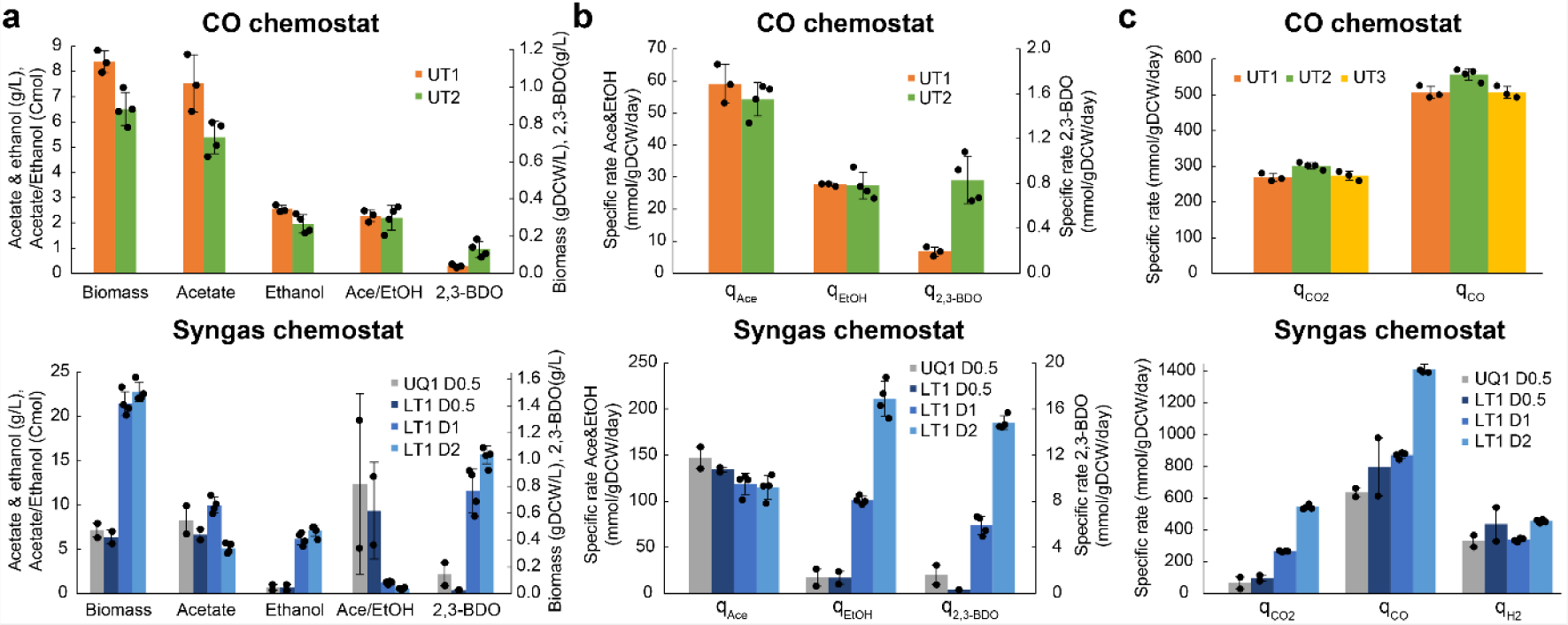
Characterisation of selected isolates from UT, UQ, and LT ALE in autotrophic chemostat cultures. (a) Biomass and by-product concentrations and acetate/ethanol (Ace/EtOH). (b) Specific by-product production rates (mmol/gDCW/day). (c) Specific gas uptake (CO, H2) and production (CO2) rates. CO chemostat data for UT1 and UT2 are from dilution rate (D) 0.5 day^−1^; for syngas chemostat data, the number following D denotes D value in day^−1^. UT3 reached steady-state only regarding gas uptake and production. Bars show average ± standard deviation between four (UT2, LT1 D1, LT1 D2), three (UT1), or two (UQ1 D0.5, LT1 D0.5) bioreplicates. See Table 1 for details. gDCW, gram of dry cell weight; 2,3-BDO, 2,3-butanediol; q, specific rate.

**Fig. 5.**
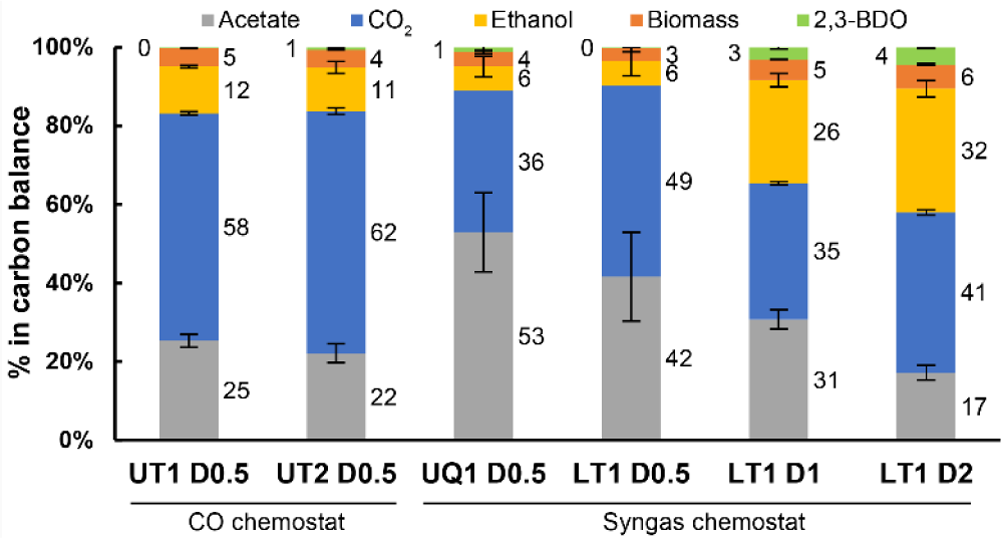
Carbon balances of selected isolates from UT, UQ, and LT ALE in autotrophic chemostat cultures. The number following D (dilution rate) denotes D value in day^−1^. Carbon flow into CO2 for UQ1 D0.5 and LT1 D0.5 were calculated by subtracting measured carbon flows into other products and biomass from 100%. Carbon recoveries for UT strains and LT D1 and D2 were normalised to 100% to have a fair comparison of carbon distributions between different conditions. Bars show average ± standard deviation between four (UT2, LT1 D1, LT1 D2), three (UT1), or two (UQ1 D0.5, LT1 D0.5) bioreplicates. See Table 1 for details. 2,3-BDO, 2,3-butanediol.

The evolved strain LT1 fulfilled all the aims of our work to create an improved model-acetogen strain available to the public: it grows without complex nutrients on minimal medium, shows high gas uptake enabling faster growth, and is robust in terms of operating continuous bioreactor gas fermentation cultures. Additionally, the metabolic feature of high carbon flux into reduced growth by-products makes LT1 an attractive chassis for metabolic engineering. We have thus deposited the LT1 strain at the DSMZ strain collection (DSM 115981) for public use and propose to make it the model-acetogen strain named *C. autoethanogenum* LAbrini (i.e. Laboratory Adapted Abrini). The improved characteristics of LAbrini hold promise for speeding up phenotyping and engineering of acetogens. For example, we currently map genotype-phenotype links for every gene in LAbrini using a systems-level approach.

### 3.4. Genetic mutations associated with improved phenotypes

We next used whole-genome DNA sequencing (WGS) to identify genetic mutations during the three ALE strategies, which may have led to different phenotypes. WGS data also allows the comparison of genetic variability between the evolved strains. We sequenced the starting strain JA1-1 and seven key evolved strains (UT1, UT2, UT3, UQ1, UQ2, UQ3, and LT1) and performed mutation analysis after high-quality genome assemblies (see Material and Methods). The starting strain for our three ALE strategies was sequenced to ensure we reliably detected genetic mutations that appeared during ALE instead of relying only on the published genome sequences for *C. autoethanogenum* JA1-1 (33, 35).

In total, 25 mutations were detected across the evolved strains, with 21 affecting protein-coding sequences (CDSs) (Fig. 6; Supplementary Table 6). 17 of the 25 mutations were single nucleotide polymorphisms (SNPs). Eight were deletions, while four being ≥ 168 bp (Fig. 6b). Two hot-spots were detected in the genome that accrued mutations during multiple ALE strategies: mutations #19‒23 around genome position 548,800 (all in the same CDS) and mutations #4‒5 around position 3,469,400 (both in the same CDS) (Fig. 6). These two genes are examples of convergent evolution under stronger autotrophic selection pressure across different ALE strategies. Principal component analysis (PCA) of all the WGS data confirmed that strains isolated from the same population (from UT or UQ ALE) are closely related (Supplementary Fig. 8). Comparison of the genomes of the superior strain LT1 and starting strain JA1-1 showed similar but not identical values: length of 4,349,910 nt, 3,942 CDSs, and 31.1% GC content for LT1; and 4,352,446 nt; 3,964 CDSs; 31.1% GC content for JA1-1.

**Fig. 6.**
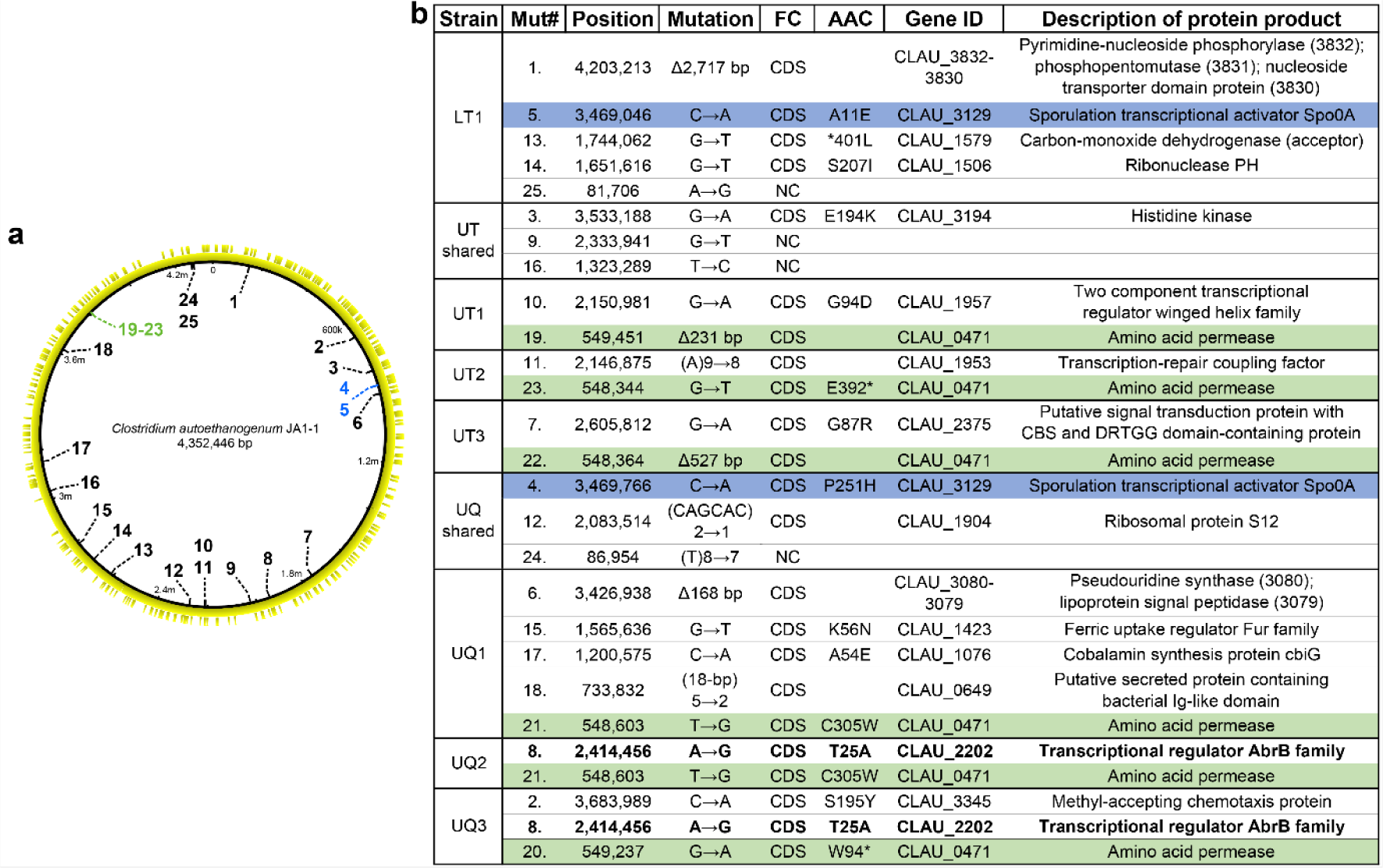
Whole-genome sequencing of the selected isolates from UT, UQ, and LT ALE and identified mutations relative to the wild-type *C. autoethanogenum* starting strain of ALE (JA1-1). (a) Distribution of the identified 25 mutations across the circular genome of JA1-1. The number denotes mutation shown in panel b (column Mut#) and colour matches the shading of mutation in panel b (exceptions are black numbers with white shading in panel b). Yellow layers are protein-coding sequences (CDS) on forward (inner) and reverse (outer) strands. (b) Details of the 25 mutations identified in the seven ALE strains. Mutations with colour shading appeared in the same genes across multiple ALE strategies, i.e. convergent mutations, while the non-convergent mutation in bold font (#8) appeared in two isolates within one ALE strategy (details shown only for the first occurrence); shading colours match colour of numbers (Mut#) on panel a. The bases within the parentheses in column “Mutation” denote the repeat sequence that decreased from the number of repeats indicated on the left of the arrow (→) to the number indicated on the right. See Supplementary Table 6 for additional details on mutations. Mut#, mutation number; FC, feature class; AAC, amino acid change; Gene ID, gene identifier; SNP, single nucleotide polymorphism; Δ, deletion; NC, non-coding; *, stop codon.

The first hot-spot at ∼548,800 position is a proposed amino acid permease (CLAU_0471; function predicted based on conserved domain PotE) that was affected by two deletions and three different SNPs: one 231 bp deletion in UT1 and one 527 bp deletion in UT3; two hits in UQ isolates and one in UT2 (mutations #19‒23 in Fig. 6b and Supplementary Table 6). Interestingly, nucleotides in the ALE starting strain JA1-1 at these three SNP positions of CLAU_0471 are fully conserved across homologs of Clostridia species (Supplementary Fig. 9). The second hot-spot at position ∼3,469,400 is the sporulation transcription factor *spo0A* (CLAU_3129) that was affected by a G-to-T SNP at two different locations in UQ and LT ALE (mutations #4‒5 in Fig. 6b and Supplementary Table 6). Similarly to the mutations in the first hot-spot (Supplementary Fig. 9), nucleotides in the starting strain JA1-1 at both positions of the second hot-spot are highly conserved across homologs of Clostridia species (Supplementary Fig. 10). UQ1 acquired a 168 bp deletion (mutation #6) across genes encoding for a lipoprotein signal peptidase (CLAU_3079) and a pseudouridine synthase (CLAU_3080) that are involved in sporulation (48) and pyrimidine metabolism, respectively (Fig. 6b and Supplementary Table 6). Interestingly, the latter gene was upregulated under butanol stress in a *spo0A* overexpression mutant strain in *Clostridium acetobutylicum* (49). Pyrimidine metabolism seems to have played an important role in improving strain phenotypes in our ALE approaches as the large 2,717 bp deletion in LT1 (mutation #1) resulted in the deletion of two genes (CLAU_3830 and 3831) and truncation of one gene (CLAU_3832) that are associated with pyrimidine metabolism (Fig. 6b and Supplementary Table 6).

Among the other spatially dispersed mutations in the genome, the mutation potentially most relevant to the improved phenotype of LT1 is a SNP (mutation #13) in the carbon monoxide dehydrogenase gene AcsA (CLAU_1579) that encodes for the CODH subunit of the CODH/acetyl-CoA synthase (ACS) protein complex (Fig. 6b and Supplementary Table 6). Mutation #13 removes the AcsA stop codon, resulting in a putative translational fusion product. Interestingly, while NTG treatment during LT ALE did not result in significantly more mutations for LT1 than what appeared during UT and UQ ALE, all but one mutation appeared in unique genes compared to UT and UQ strains (Fig. 6b). The only CDS mutation (#3) present in all UT ALE strains was found in a predicted histidine kinase (CLAU_3194), which is interesting as some histidine kinases regulate sporulation and metabolism in Clostridia (50). Notably, one SNP appeared during UQ ALE in a usually highly conserved position in Clostridia species (Supplementary Fig. 11) - mutation #8 in an AbrB family transcriptional regulator (CLAU_2202) (Fig. 6b and Supplementary Table 6). Interestingly, CLAU_2202 is among the top 100 abundant proteins in *C. autoethanogenum* (51). Other potentially relevant mutations in CDSs appeared in a ribosomal protein S12 (mutation #12), and transcription-associated genes (mutations #10, #11, and #15). A homolog of the Ferric uptake regulator of Fur family (CLAU_1423) that acquired mutation #15 in our work – the putative transcriptional regulator PerR – has acquired numerous mutations within the ALE experiments compiled in the ALEdb database (52). Future advances in quantifying genotype-to-phenotype links in acetogens will assist in interpreting the genetic changes we detected and improve experimental evidence-based (vs. computationally derived) annotation of *C. autoethanogenum* genome.

### 3.5. Reverse genetic engineering recovers superior features of our ALE strains

The causality of mutations observed during ALE can be evaluated by reconstructing the genetic modifications in the starting strain of ALE, i.e. using reverse genetic engineering. Observation of convergent evolution by appearance of mutations in *spo0A* during both LT and UQ ALE (Fig. 6b) is likely significant due to the role of *spo0A* as a master regulator of sporulation and solventogenesis in Clostridia (53–56). Interestingly, mutation #10 observed during UT ALE (in UT1) occurred in a two-component transcriptional regulator CLAU_1957 (based on GenBank CP012395.1 annotation; (33)) that could be potentially involved in sporulation processes (e.g. phosphorelay, sporulation activator; (57)) based on predictions using protein sequence or structure homology by DeepFRI (58), ProteinInfer (59), and UNIPROT database (60) (Supplementary Table 7; UniProtKB U5RXC5). Furthermore, this mutation is predicted to have occurred in a response regulator receiver domain (one of the two predicted domains) (61). Therefore, we targeted to mutate *spo0A* or CLAU_1957 in our ALE starting strain JA1-1 to investigate whether alterations to cellular sporulation processes could, at least partially, replicate the improved phenotypes obtained during our ALE: faster growth, growth without complex nutrients, and robustness in continuous culture operation.

We used CRISPR/Cas9-aided homologous recombination (28) for reverse genetic engineering of JA1-1. Mutations #4 and #5 occurred across LT and UQ ALE (Fig. 6b) in the linker domain between the activator domains and in the receiver domain of Spo0A, respectively (62). Mutations in the same domains in other Clostridia have led to the same phenotype as *spo0A* deletion, i.e. non-sporulating cells (63–65). We thus aimed to investigate the effects of both mutations #4 and #5 through *spo0A* deletion. Both of the two sgRNAs we used yielded 100% gene deletion efficiency as *spo0A* was deleted in all screened colonies (Supplementary Fig. 12 and Supplementary File 4 and 5); this strain was named RE1. Mutation #10 was a SNP in CLAU_1957, and we identified only one potential sgRNA that would realize the G-to-A mutation and disrupt the PAM sequence, thus preventing Cas9 from cleaving the target site again after successful homologous recombination. This sgRNA yielded 100% gene-editing efficiency as all screened colonies showed the desired SNP (Supplementary File 6 and 7); this strain was named RE2. Strikingly, RE1 autotrophic batch bottle cultures without YE showed >4-fold faster growth than JA1-1 with YE and similar values on both CO and syngas as LT1 (Fig. 7a and Fig. 2). We cultured RE2 on CO, i.e. same gas feed as used for UT1 characterization (ALE strain with same SNP as RE2), and also this reverse engineered strain displayed significantly faster growth (>3-fold) without YE compared to JA1-1 with YE and similar μ_max_ to UT1 (Fig. 7a and Fig. 2).

**Fig. 7.**
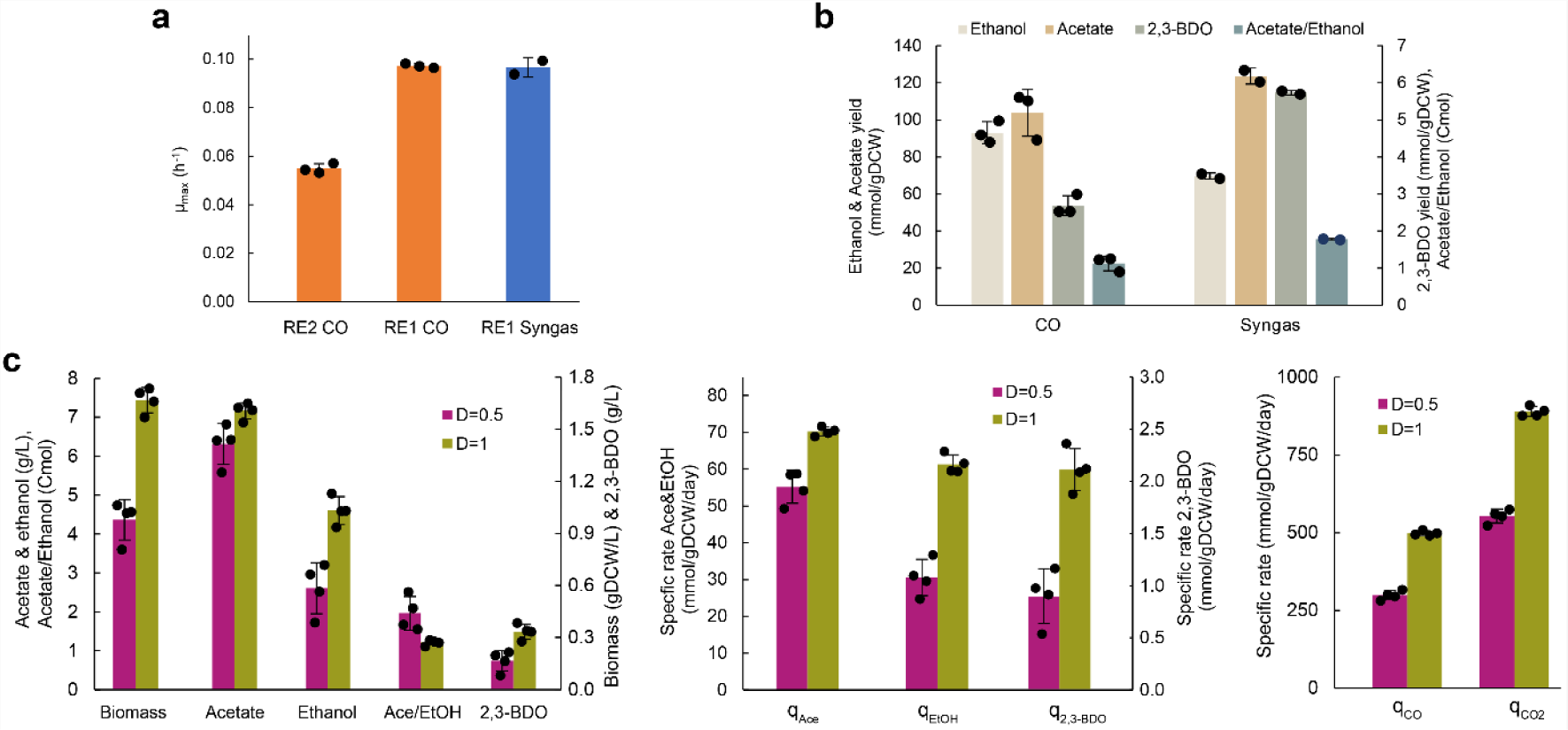
Autotrophic characterization of reverse engineered strains RE1 and RE2 in batch and chemostat cultures. (a) Maximum specific growth rate (μmax) in bottle batch cultures with yeast extract (YE)-free medium. (b) Production yields (mmol/gDCW) of growth by-products and acetate/ethanol (Ace/EtOH) of strain RE1 in bottle batch cultures. (c) RE2 growth characteristics in CO chemostats at dilution rate (D) 0.5 and 1 day^−1^. See Fig. 4 legend for parameter details. Bars show average ± standard deviation between four (panel c), three (RE2 CO, RE1 CO, and CO data on panel b), or two (RE1 Syngas and Syngas data on panel b) bioreplicates. gDCW, gram of dry cell weight; 2,3-BDO, 2,3-butanediol; q, specific rate.

Quantification of by-product patterns of RE1 in autotrophic bottle cultures (Fig. 7b) showed similar yields to JA1-1 on CO (see Fig. 2c; growth with YE) and higher acetate/ethanol ratios compared to LT1 (see Fig. 2c and d) while ethanol and acetate production did not coincide in RE1. The performance of RE2 was tested in bioreactor continuous cultures as we experienced significant difficulties with growing UT strains to high biomass concentrations on CO (see section 3.3 above). We efficiently operated the culture at D = 0.5 day^−1^ and achieved steady-state at high gas uptake rates and biomass concentrations at D = 1 day^−1^ (double of UT and UQ strains; see Table 1) (Fig. 7c). The robustness of RE2 in continuous cultures was surprising since just one SNP was introduced into the JA1-1 background. Despite the significant difference in bioreactor operation between UT1 and RE2 cultures, the two strains displayed somewhat similar growth characteristics (e.g. biomass concentration, specific by-product production rates, specific gas rates) at D = 0.5 day^−1^ except >4-fold higher 2,3-BDO production for RE2 (Fig. 4 and 7c). Comparison of RE2 performance at D = 1 day^−1^ is only possible with LT1 data (Table 1). RE2 showed lower production of acetate and ethanol (Fig. 4 and 7c), though LT1 cultures were fed with syngas (RE2 with CO). This is also reflected in the carbon balance where significantly more carbon is diverted to CO_2_ for RE2 (Supplementary Fig. 13 and Fig. 5), which is consistent with another strain of *C. autoethanogenum* diverting more carbon to CO_2_ on CO compared to syngas (9). Higher D led to similar effects for RE2 and LT1 (Fig. 4 and 7c). In conclusion, reconstructing mutations in two sporulation-related genes was sufficient to recover all three superior features of our ALE strains.

We tested next if the reverse engineered mutations in sporulation-related genes might have affected sporulation in the strains. For this, we cultured RE1, RE2, and JA1-1 both autotrophically and heterotrophically and used microscopy to detect potential differences in sporulation from stationary phase samples. We also plated heterotrophic cultures before and after a heat-shock with the aim to observe potential heat-resistant spores. Neither microscopy (Supplementary Fig. 14 and 15) nor plating (Supplementary Fig. 16) identified any spores or sporulating cells for any of the strains (including JA1-1). Although the authors who isolated *C. autoethanogenum* JA1-1 report it as a spore-former (27), a recent work also failed to detect JA1-1 forming spores (66). The latter and our observations are supported by a review on Clostridial sporulation concluding that sporulation is rarely observed in acetogens (62).

Mutations in sporulation-related genes have led to significant effects on Clostridial phenotypes other than sporulation (62). We thus profiled the proteomic response of JA1-1 to the reverse engineered mutations as functional genomics has been informative for interpreting genotype-phenotype links in engineered strains. For this, JA1-1, RE1, and RE2 were grown in syngas bottle cultures with YE in the media as JA1-1 does not grow without YE. We note that these proteome expression patterns are affected by variable μ_max_ values (Fig. 8a) and potential differences in consumption of YE components between the strains. Our analysis identified 284 differentially expressed proteins (fold-change > 2 and q-value < 0.05) between RE1 and JA1-1, and 514 between RE2 and JA1-1 (Supplementary Table 5). Proteomic responses to either mutation were similar in general (Fig. 8b; Supplementary Fig. 17) and shared 205 differentially expressed proteins potentially due to the 66-fold down-regulation of Spo0A in RE2 (Fig. 8c; Supplementary Table 5). Notably, peptides for five sporulation-related proteins highly expressed in JA1-1 (CLAU_1942, 1946, 3117, 3233, 3248) were not detected in either engineered strain (also not for Spo0A in RE1). At the same time, three histidine kinases and two phosphatases potentially linked to Spo0A regulation were up-regulated in both RE1 and RE2 (Fig. 8c), including CLAU_3194 that was mutated in all UT isolates (mutation #3 in Fig. 6b). Phosphate metabolism was further affected by 12 differentially expressed kinases not shared between the two strains (Supplementary Table 5).

**Fig. 8.**
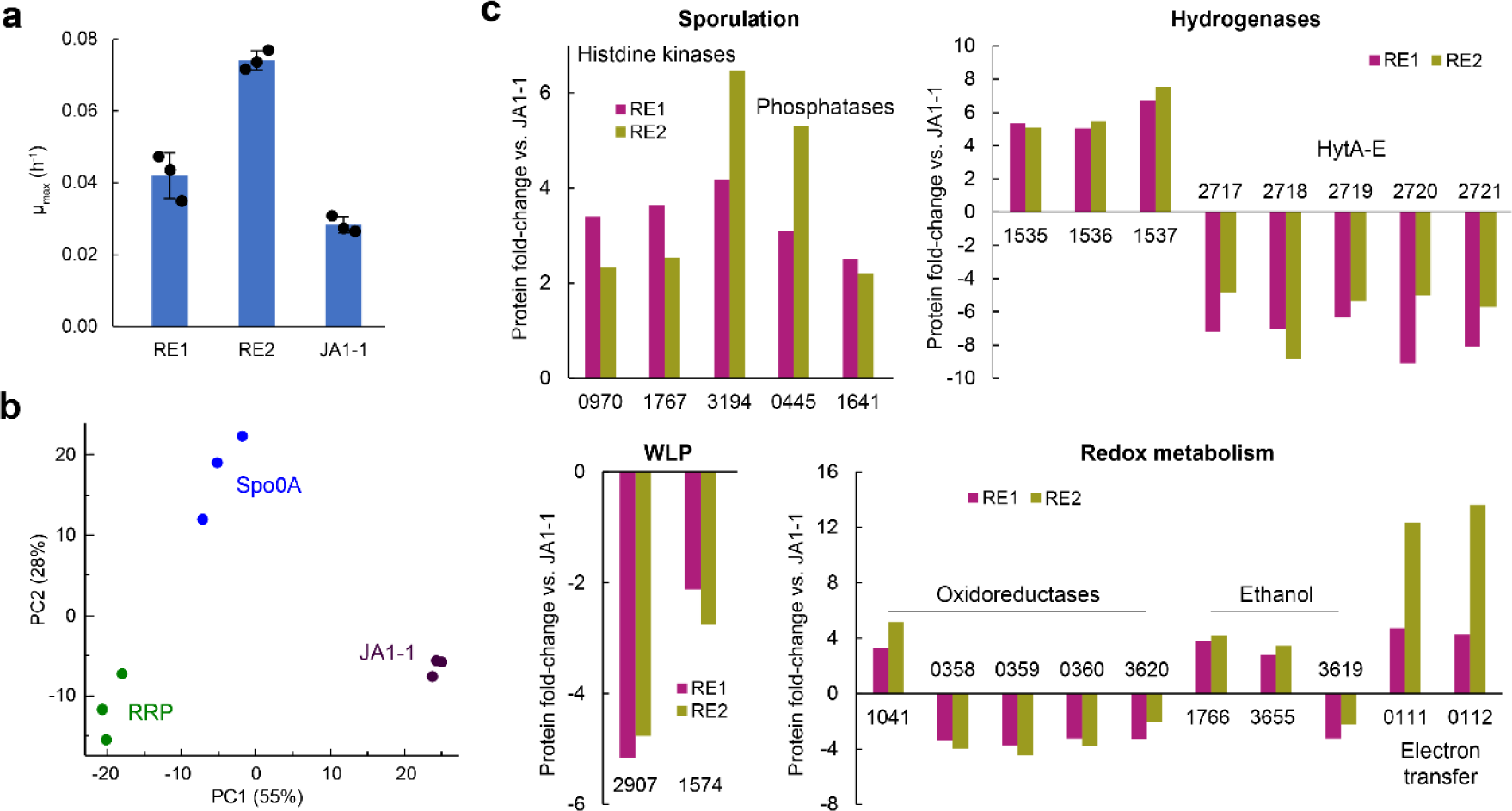
Proteomic responses of the wild-type *C. autoethanogenum* starting strain of ALE (JA1-1) to the reversed engineered mutations in syngas bottle cultures. (a) Maximum specific growth rate (μmax) for reverse engineered strains RE1, RE2, and JA1-1 with yeast extract (YE) in medium. Bars show average ± standard deviation between three bioreplicates. (b) Principal component analysis (PCA) of protein MS intensities for the three bioreplicate cultures of RE1, RE2, and JA1-1. PC, principal component. (c) Potential key proteins differentially expressed (fold-change > 2 and q-value < 0.05) for both RE1 vs JA1-1 and RE2 vs JA1-1 comparisons. Protein IDs are preceded with CLAU_. See Supplementary Table 5 for data.

Regarding functionalities specifically relevant for acetogens, repression of the electron-bifurcating hydrogenase HytA-E complex was potentially compensated by induction of hydrogenase CLAU_1535 and its associated proteins in both strains (Fig. 8c). In addition, a formate dehydrogenase (CLAU_2907) and the methylene-THF dehydrogenase (CLAU_1574) in the WLP were down-regulated in both strains. Also redox metabolism was markedly affected by both mutations as oxidoreductases, proteins involved in ethanol metabolism and electron transfer were differentially expressed in both strains compared to JA1-1 (Fig. 8c). The necessary transcriptional changes needed to recover the superior phenotypes of our ALE strains through reverse engineering could potentially have been realised through 2-to-10-fold up-regulation of seven and/or 2-to-6-fold down-regulation of two transcriptional regulators shared for both strains (Supplementary Table 5). Mutations in either sporulation-related gene thus lead to significant proteomic rearrangements in *C. autoethanogenum*.

## 4. DISCUSSION

While gas fermentation using *C. autoethanogenum* has been deployed commercially for ethanol production (7), non-commercial acetogen strains struggle to grow on minimal medium, show slow gas uptake and growth, and are challenging to run in continuous bioreactor cultures. To overcome these limitations, we used three different and independent ALE strategies in this work to obtain strains with improved phenotypes of which the strain named LAbrini (LT1) showed superior performance in acetogen gas fermentation. We characterised the evolved strains using bottle and bioreactor cultures, gas analysis, metabolomics, transcriptomics, and DNA sequencing. Finally, reverse genetic engineering of sporulation-related mutations recovered all three superior features of our ALE strains.

During one of the ALEs (UT), we noticed a declining μ_max_ during the last ⅓ of the ALE (Fig. 1b). This implies that evolutionary pressures other than towards growing faster were involved in this ALE approach of serial batch culture propagation. Similar trends have been seen before (45, 46). Additionally, UT ALE populations showed a declining trend of acetate/ethanol with faster growth (Fig. 1c). Some isolates of the ALE populations of this work displayed faster growth coupled with higher acetate production (Supplementary Fig. 5). These observations at isolate and population levels suggest that cells can find various metabolic routes across different ALE approaches to achieve faster growth, as noted before (67, 68). We hypothesise that cross-feeding between subpopulations took place similar to previous bacterial ALEs (69–72). Future co-culturing experiments of isolates with different phenotypes (e.g. high and low acetate producers) might shed light on these possibilities.

Characterisation of evolved strains in batch and chemostat cultures revealed that batch bottle cultures might fail to predict performance in continuous cultures. For example, UT1 and UT2 showed nearly identical q_EtOH_ and acetate/ethanol ratios in chemostats after very different results in batch cultures and only LT1 could thrive above D = 0.5 day^−1^ while most evolved strains showed similar μ_max_ in batch cultures, which was > 0.5 day^−1^. Furthermore, efforts to operate UT1 chemostats at higher D and gas-liquid mass transfer seemed to have been compromised by CO toxicity (see below), which was not present in batch cultures with lower gas-liquid mass transfer. These results emphasise the importance of testing novel strains (e.g. acquired using ALE or metabolic engineering) in industrially-relevant continuous fermentation conditions (7). We also note that the isolation of LT1 from a chemostat culture after NTG mutagenesis could have contributed towards LT1 superior performance in our bioreactor continuous cultures. Thus, isolating evolved strains from operational conditions similar to those where the strain would eventually be used for scientific investigation or industrial exploitation could be valuable. When comparing our different ALE approaches, the combination of NTG with selective pressure during continuous cultivation in LT ALE proved to be the most successful strategy. NTG treatment presumably introduced high genetic diversity, setting the stage for subsequent selection under operational conditions. Indeed, LT1 was superior to UQ ALE isolates that were also obtained from chemostats but without prior treatment with a mutagen. These results support the effectiveness of NTG-induced mutagenesis combined with selective pressure in microbes (73, 74).

Transcriptome analysis of the potential “CO toxicity” state of UT1 chemostat at D = 1 day^−1^ revealed a distinct transcriptional profile compared to other cultures, including UT1 in a “non-CO toxicity” but unstable culture at D = 1 day^−1^. UT1 in the “CO toxicity” state showed strong up-regulation of numerous hypothetical and sporulation-related proteins (Supplementary Table 2). These proteins serve as promising candidates for potential overexpression to play a role in mitigating CO toxicity. Intriguingly, we also observed ∼2-fold down-regulation of components of the HytA-E/formate dehydrogenase (FdhA) complex (75), which plays a key role in CO_2_ reduction to formate in *C. autoethanogenum* using NADPH and reduced ferredoxin (Fd) or H_2_ (29, 75, 76). Wang and co-workers who purified the complex hypothesised that it might also protect cells from over-reduction of NADP and Fd pools during growth on CO (75). They propose that lowered CO oxidation increases the CO concentration in cells that inhibits CO_2_ reduction to formate by the complex that lowers reoxidation of Fd. The solution for avoiding this over-reduction state is NADPH and Fd reoxidation by HytA-E through proton reduction to H_2_. Our results are consistent with their hypothesis as we detected H_2_ production when trying to improve COUR of the UT1 “CO toxicity” culture by decreasing gas-liquid mass transfer, i.e. CO oxidation (Supplementary Fig. 2). We thus hypothesise that such regulation could be linked to the down-regulation of HytA-E/FdhA complex component genes. The metabolism of *C. autoethanogenum* thus displays reactional plasticity in choosing electron sinks to drive CO_2_ reduction as proposed before (75, 77). Such flexibility would realise growth advantages under changing environmental conditions.

Our ALE experimental data analysis confirmed previous observations that different ALE approaches often lead to similarly improved phenotypes but through various underlying genetic changes (67–69, 71, 72). Interestingly, a proposed amino acid permease (CLAU_0471) was affected in all the isolated strains in two different ALE approaches (UT and UQ) by two distinct deletions and three SNPs. It is unclear why a proposed amino acid permease was under such strong selective pressure when the complex nutrient YE was removed from the medium. The other gene that was mutated during two of the ALEs (LT and UQ) was *spo0A* (CLAU_3129) which is considered a master regulator of sporulation and solventogenesis in Clostridia (53–56). Down-regulation or inactivation of *spo0A* in *C. acetobutylicum* causes a reduction in sporulation and increased solvent production (78, 79). However, in other Clostridia Spo0A can improve fitness (65) and regulate metabolic and virulence factors outside the sporulation process (80). Interestingly, mutation #10 in a predicted phosphorelay regulator (CLAU_1957) and mutation #3 in a proposed histidine kinase (CLAU_3194) could be linked to affect sporulation in UT1 as latter regulators are involved in phosphorylation circuits involving histidine kinases (81) and as similar kinases regulate sporulation and solventogenesis in *Clostridium beijerinckii* and *C. acetobutylicum* (50, 82). Additionally, both histidine kinases (53, 54) and CLAU_1957 (83) could affect Spo0A directly. Mutations in two-component sensor systems composed of histidine kinases and response regulators that appeared in ALE experiments with other bacteria were postulated to be related to general adaptation to the growth environment (84). Sporulation could have also been affected during UQ ALE (in UQ1) by truncation of a lipoprotein signal peptidase (CLAU_3079), which is involved in sporulation of *Clostridium difficile* (48) and deletion of the gene pseudouridine synthase (CLAU_3080) that was upregulated in a *spo0A* overexpression mutant strain in *C. acetobutylicum* (49). Notably, protein expression of CLAU_3080 was up-regulated 3-fold in our RE2 strain (Supplementary Table 5). Both reverse engineered mutations triggered widespread proteomic rearrangements demonstrating their effects beyond the sporulation process. Similarly, altered expression of genes in Clostridial sporulation networks regulate gene expression across various metabolic functions (54, 62). We hypothesise that mutations in the above-described genes could have emerged from selective pressure towards faster growth since suppressing sporulation functionalities would save cells valuable energy to grow faster, for instance, through conserving a phosphate group (54). This is consistent with our reverse engineering results of strains with mutations in *spo0A* or CLAU_1957 growing significantly faster than wild-type *C. autoethanogenum*. The specific regulatory mechanisms governing the proteome changes that realised the superior phenotypes remain to be elucidated.

The CODH/ACS complex realises CO oxidation/CO_2_ reduction and acetyl-CoA synthesis (85), and its activity is essential for the autotrophic growth of *C. autoethanogenum* (86). The *acsA* (CLAU_1579) stop codon removal in strain LT1 has been reported in an independent study that observed translational readthrough at the protein level (86). The *C. autoethanogenum* AcsA protein is uniquely truncated compared to other acetogens. The other potential key genetic modification in LT1 seems to be associated with pyrimidine metabolism evidenced by the large 2,717 bp deletion affecting three genes in pyrimidine metabolism. This could realise a faster supply of RNA monomers, allowing higher translation rates and faster growth. Thus, the superior performance of LAbrini in continuous bioreactor cultures and its ability to grow in the absence of YE is due to its genotype and the selection strategy employed in its development. While growth without YE has been reported for acetogens before (25, 26), it is unclear if these phenotypes resulted from physiological adaptation or evolution. Further studies are required to link mutations to its phenotype, key structural/functional attributes of the ACS/CODH enzyme complex, and regulation.

## CONCLUSIONS

Our work is important for two reasons. First, we offer a superior model-acetogen strain – LAbrini – for the community that grows fast without complex nutrients, is robust for continuous gas fermentation cultures, and shows high carbon flux to reduced by-products. These strain characteristics will speed up both phenotyping and metabolic engineering of acetogens. Second, our comprehensive ALE approach with three strategies offers insights into convergent evolution trajectories in acetogens and highlights novel engineering targets for improved phenotypes. Further work is needed to validate the causality of other mutations uncovered in our ALEs by reconstructing mutations of interest, both in combination and in isolation, and assaying their contribution to observed phenotypes. Future ALE experiments with LAbrini should potentially be designed to select for improved H_2_ uptake, CO tolerance, and by-product tolerance.

## ACKNOWLEDGEMENTS

This work was funded by the European Union’s Horizon 2020 research and innovation programme under grant agreement N810755 and the Estonian Research Council’s grant agreement PSG289. Australian Government funding through its investment agency, the Australian Research Council, towards the ARC Centre of Excellence in Synthetic Biology (CE200100029) is gratefully acknowledged. We acknowledge support from the Australian Centre for Ecogenomics for Whole Genome Sequencing of UQ strains. We thank Kristina Reinmets and Ugochi Jennifer Nwaokorie for their valuable comments on the manuscript, Shilpa Nagaraju for help with Supplementary Table 1, and Nastassia Shtaida for help with microscopy.

## DATA AVAILABILITY STATEMENT

RNA sequencing data have been deposited in the NCBI Gene Expression Omnibus repository under accession number GSE225123. Proteomics data have been deposited to the ProteomeXchange Consortium (http://proteomecentral.proteomexchange.org) via the PRIDE partner repository (44) with the data set identifier PXD047330.The genome assembly of strain LT1 (“LAbrini”) has been deposited to the NCBI genome submission portal with reference Genbank CP110420.

## AUTHOR CONTRIBUTIONS

Conceptualization: HI, JKH, AH, EM, SB and KV; Methodology: HI, JKH, AH, SB, KMS, AYS, MJP, LAL, KRM, RAGG, GH, JD, LT, HZ, ROJ, VR, HS, JJ, MK, EM, and KV; Formal analysis: HI, JKH, AYS, MJP, LAL, KRM, SB, VR, HS, JJ, and KV; Investigation: HI, JKH, AH, SB, KMS, AYS, MJP, LAL, KRM, RAGG, GH, JD, LT, HZ, ROJ, and KV; Resources: SDS, MK, EM, and KV; Writing – Original Draft: HI and KV; Writing – Review & Editing: HI, JKH, AH, SB, KMS, AYS, MJP, LAL, KRM, RAGG, LT, HZ, ROJ, VR, JJ, SDS, MK, EM, and KV; Supervision: MK, EM, and KV; Project Administration: MK, EM, and KV; Funding Acquisition: SDS, MK, EM, and KV.

