## Supplementary Figures for "Autotrophic adaptive laboratory evolution of the acetogen *Clostridium autoethanogenum* delivers the gas-fermenting strain LAbrini with superior growth, products, and robustness"

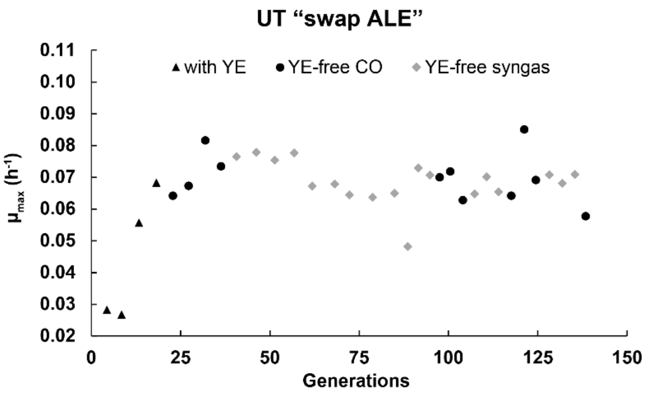


**Supplementary Fig. 1.** Change of maximum specific growth rate ($\mu_{max}$) with batch culture generations during UT "swap ALE" (see text for details). YE, yeast extract.

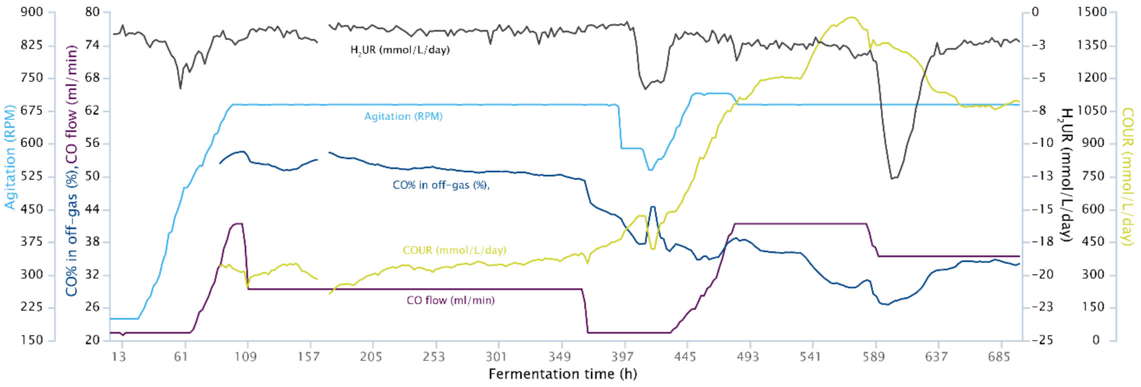


**Supplementary Fig. 2.** Chemostat fermentation profile of UT2 at a dilution rate of 1 $day^{-1}$ on CO. Data until ~380 h is of the "CO toxicity" steady-state (in Table 1) and from ~500 h of the non-stable "non-CO toxicity" state (see main text for details). Note the increase of the CO uptake rate (COUR; mmol/L/day) after gas-liquid mass transfer adjustment between ~380 to ~440 h. Off-gas analysis was unavailable between ~162 to ~175 h due to technical problems. RPM, rounds per minute; $H_2UR$, $H_2$ uptake rate (mmol/L/day).

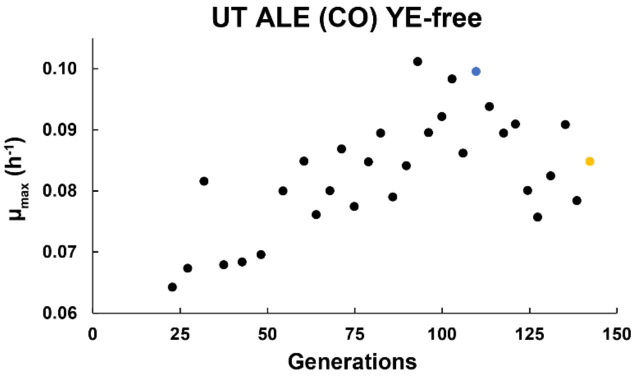


**Supplementary Fig. 3.** Change of maximum specific growth rate ($\mu_{max}$) with batch culture generations on YE-free medium during UT ALE. Data are same as on Fig. 1b. Blue and yellow circles denote cultures that were used for isolating single colonies (see text for details). YE, yeast extract.

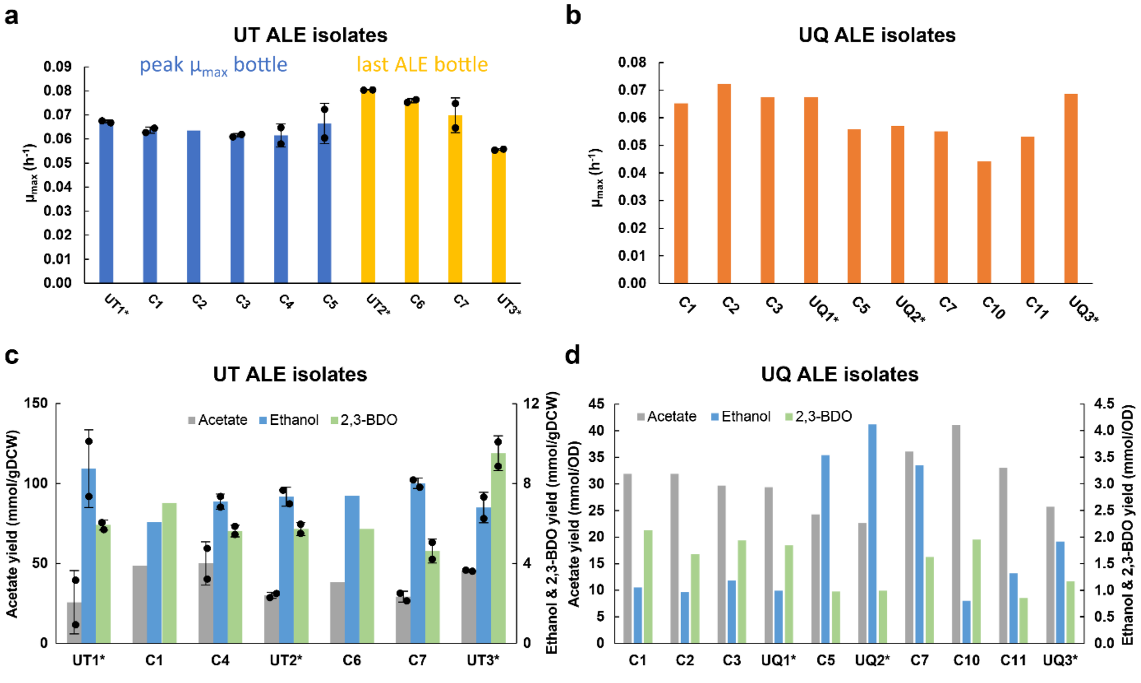


**Supplementary Fig. 4.** Preliminary screen of growth and by-product patterns of the 20 isolates from UT and UQ ALE in batch cultures. (a) Maximum specific growth rate ($\mu_{max}$) of UT ALE isolates. Bar colour matches the marker colour of culture of UT ALE on Fig. 1b from which the colonies were isolated. Bars show average ± standard deviation between two bioreplicates, except for C2 of a single batch culture. (b) $\mu_{max}$ of UQ ALE isolates. (c) Production yields (mmol/gDCW) of growth by-products of UT ALE isolates. Bars show average ± standard deviation between two bioreplicates, except for C1 and C6 of a single batch culture. (d) Production yields of growth by-products of UQ ALE isolates. UT and UQ data from growth on CO and syngas, respectively. Asterisk denotes isolate selected for further detailed characterisation (data on Fig. 2). For UQ ALE isolates, bars show data of a single batch culture. gDCW, gram of dry cell weight; 2,3-BDO, 2,3-butanediol.

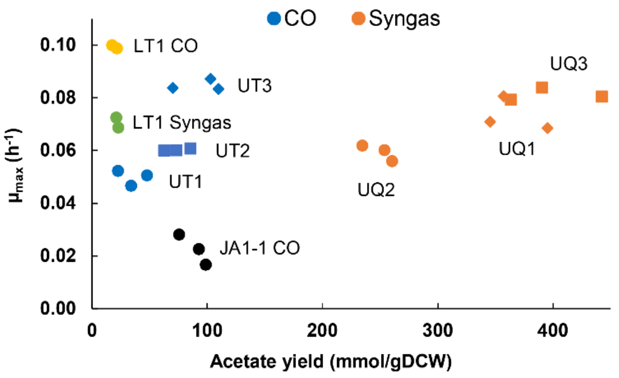


**Supplementary Fig. 5.** Correlation between maximum specific growth rate ($\mu_{max}$) and acetate production yield (mmol/gDCW) in autotrophic batch cultures of selected isolates from UT, UQ, and LT ALE, and the wild-type *C. autoethanogenum* starting strain of ALE (JA1-1). Data for UT, UQ, and LT isolates are from yeast extract (YE)-free medium, JA1-1 data are from medium with YE. gDCW, gram of dry cell weight.

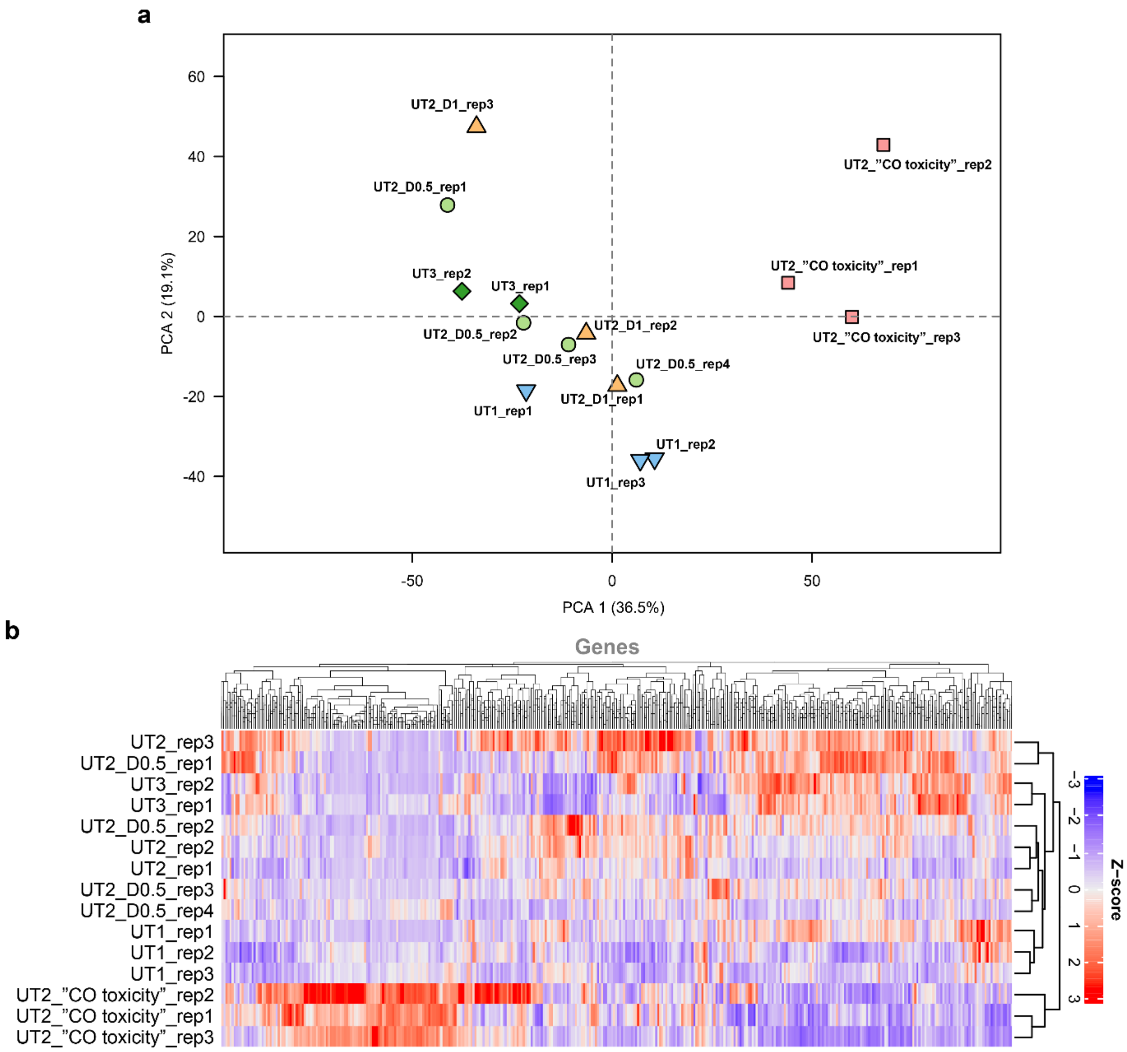


**Supplementary Fig. 6.** Transcriptional profiles of UT isolates in autotrophic chemostat cultures. (a) Principal component analysis (PCA) of transcript abundances (RPKM). (b) Hierarchical clustering of transcript abundances (Z-scores based on RPKMs). The number following dilution rate (D) denotes D value in day$^{-1}$. See Table 1 for more details. PC, principal component; rep, bioreplicate.

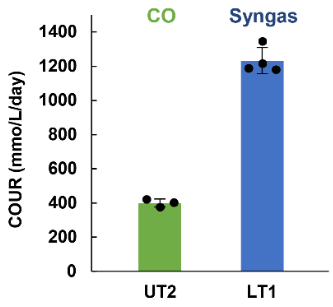


**Supplementary Fig. 7.** Comparison of CO uptake rates (COUR; mmol/L/day) of UT2 and LT1 isolates in chemostats at a dilution rate of 1 $day^{-1}$. Bars show average ± standard deviation between three (UT2) or four (LT1) bioreplicates. See Table 1 for details.

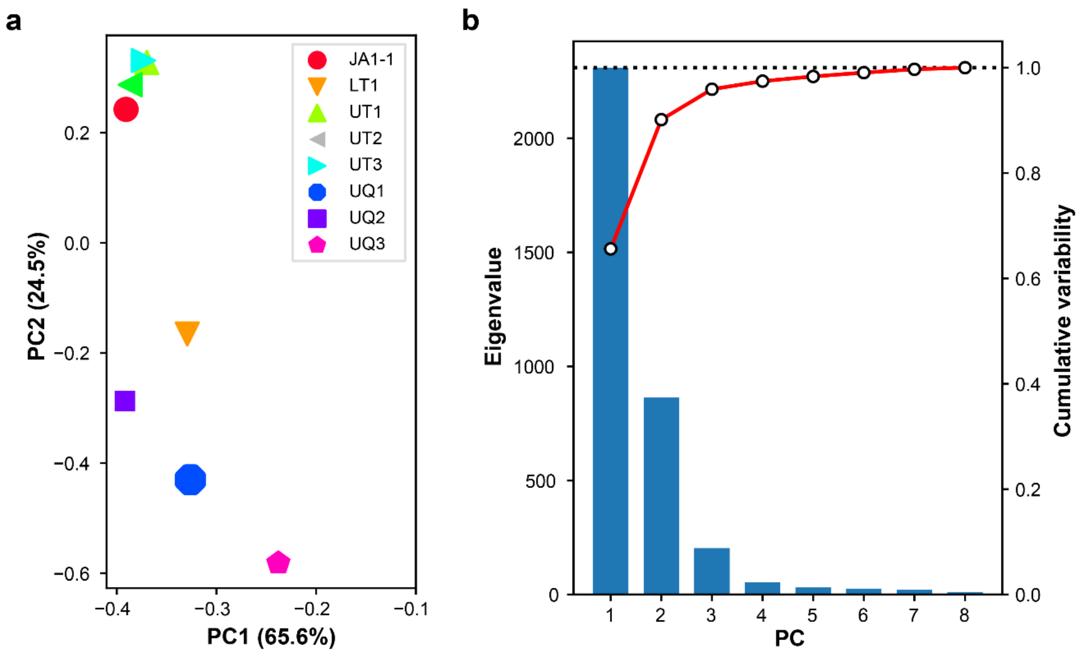


**Supplementary Fig. 8.** Variability between genome sequences of the seven evolved strains and the starting strain of ALE (JA1-1). (a) Principal component analysis (PCA) of sequence variability. (b) Scree plot of PCA. PC, principal component.

### Mutation #19

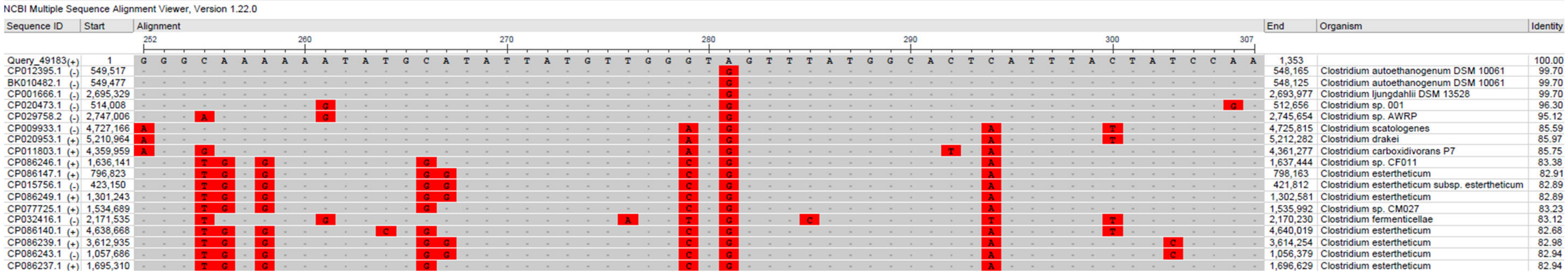


### Mutation #20

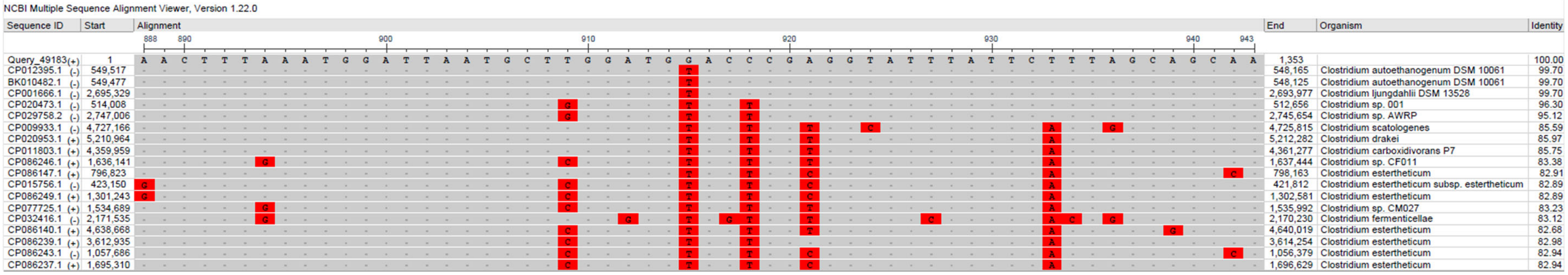


### Mutation #22

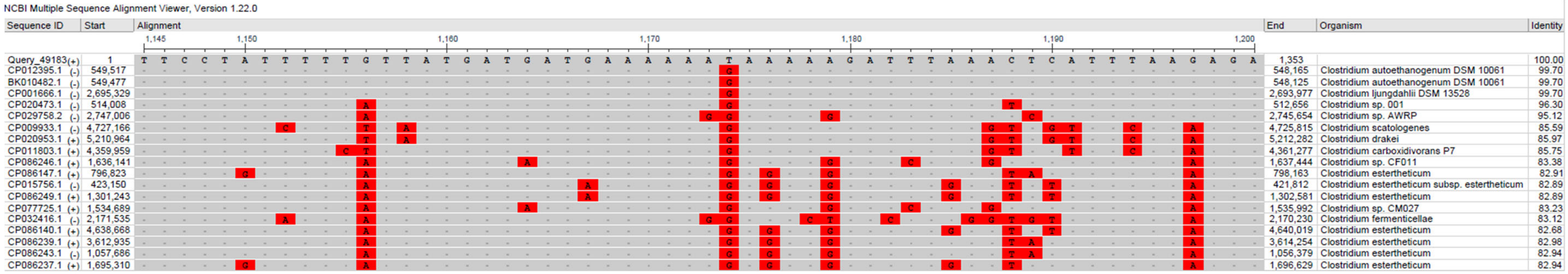


**Supplementary Fig. 9.** Conservation of mutations #19, 20, and 22 across homologs of Clostridia species. Positions of mutations: #19 at 281 nt; #20 at 915 nt; #22 at 1,174 nt. Obtained using NCBI blastn suite against nucleotide collection (nr/nt) with default settings, 80% identity cut-off, and search optimised for highly similar sequences (megablast).

### Mutation #5

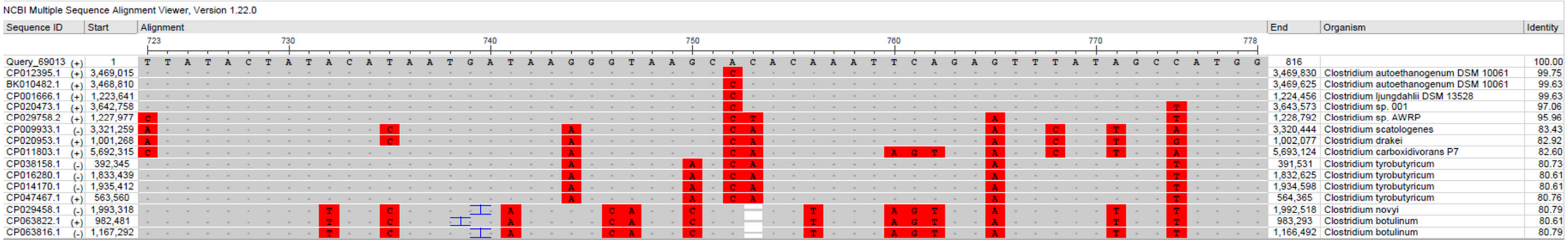


### Mutation #6

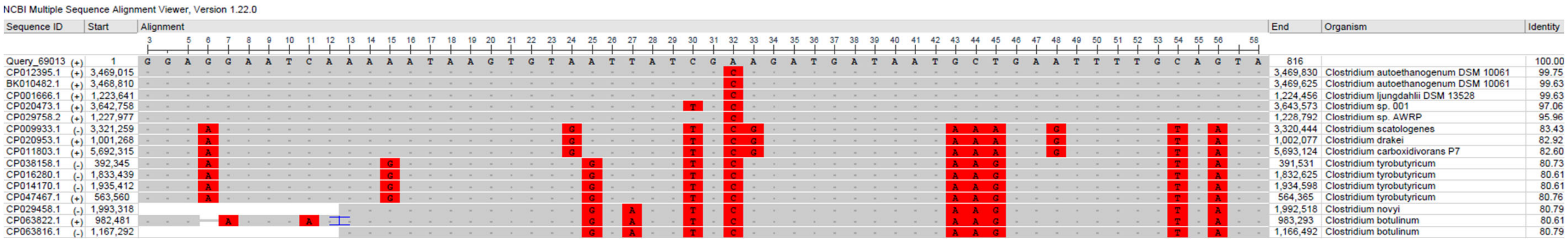


**Supplementary Fig. 10.** Conservation of mutations #5–6 across homologs of Clostridia species. Positions of mutations: #5 at 752 nt; #6 at 32 nt. Obtained using NCBI blastn suite against nucleotide collection (nr/nt) with default settings, 80% identity cut-off, and search optimised for highly similar sequences (megablast).

### Mutation #8

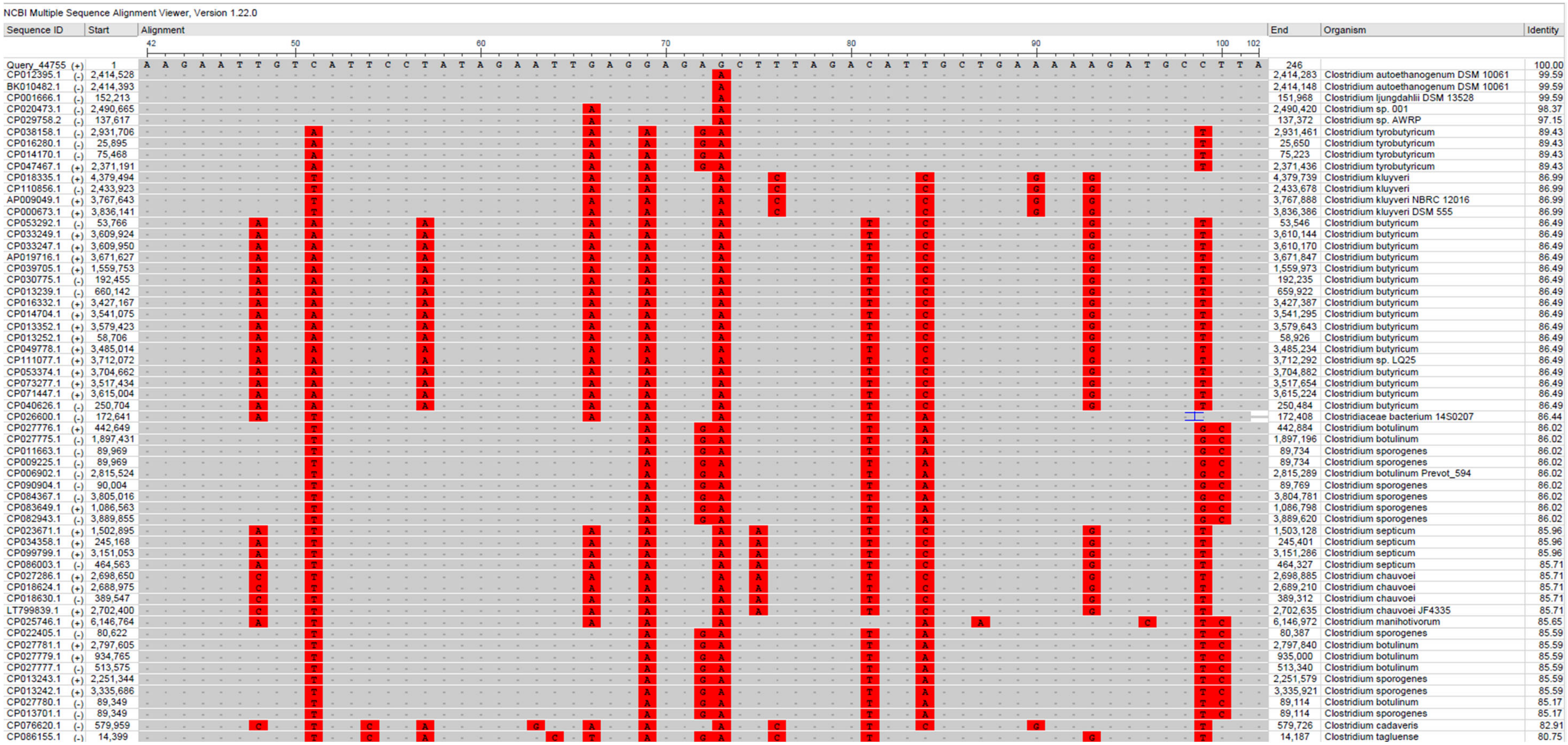


**Supplementary Fig. 11.** Conservation of mutation #8 across homologs of Clostridia species. Position of mutation #8 at 73nt. Obtained using NCBI blastn suite against nucleotide collection (nr/nt) with default settings, 80% identity cut-off, and search optimised for highly similar sequences (megablast).

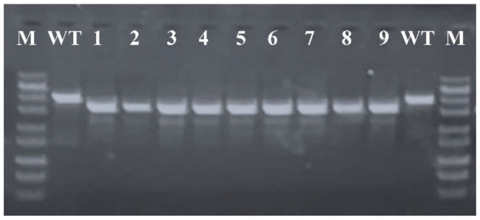


**Supplementary Fig. 12.** PCR screening of transformant colonies for *spo0A* deletion. M, DNA ladder; WT, starting wild-type strain for ALE in this work (JA1-1); 1–9, nine screened transformant colonies.

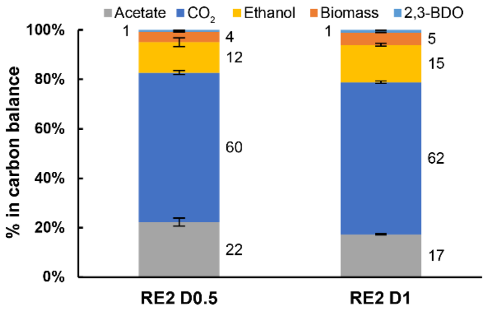


**Supplementary Fig. 13.** Carbon balances of RE2 autotrophic chemostat cultures. The number following D (dilution rate) denotes D value in day$^{-1}$. Carbon recoveries were normalised to 100% to have a fair comparison of carbon distributions between conditions and with data of ALE cultures (Fig. 5). Bars show average ± standard deviation between four bioreplicates (see Table 1). See Table 1 for details. 2,3-BDO, 2,3-butanediol.

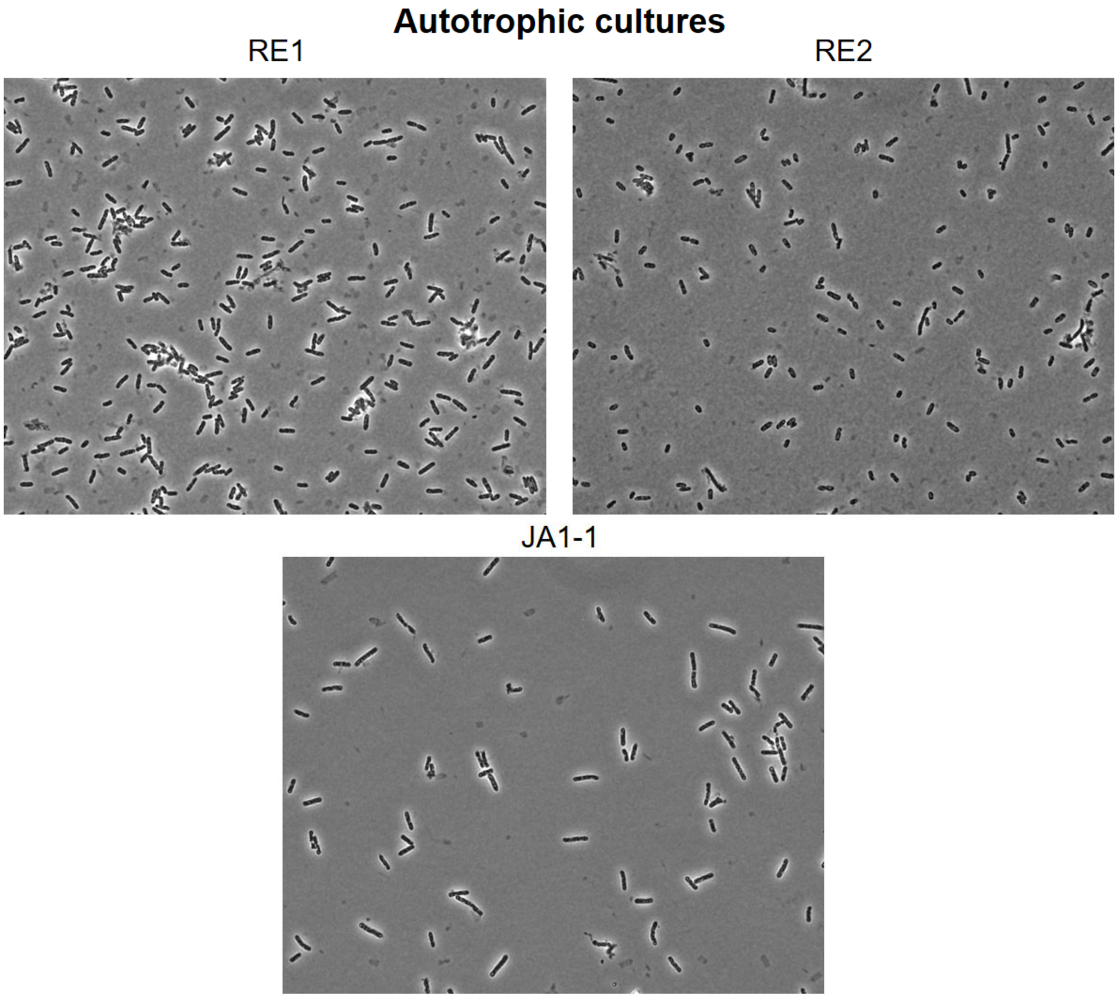


**Supplementary Fig. 14.** Micrographs showing lack of spores or sporulating cells in autotrophic cultures of JA1-1, RE1, and RE2 after ~40 days of incubation. Micrographs taken regularly during the incubation period showed same results. Spores or endospores are expected to be white.

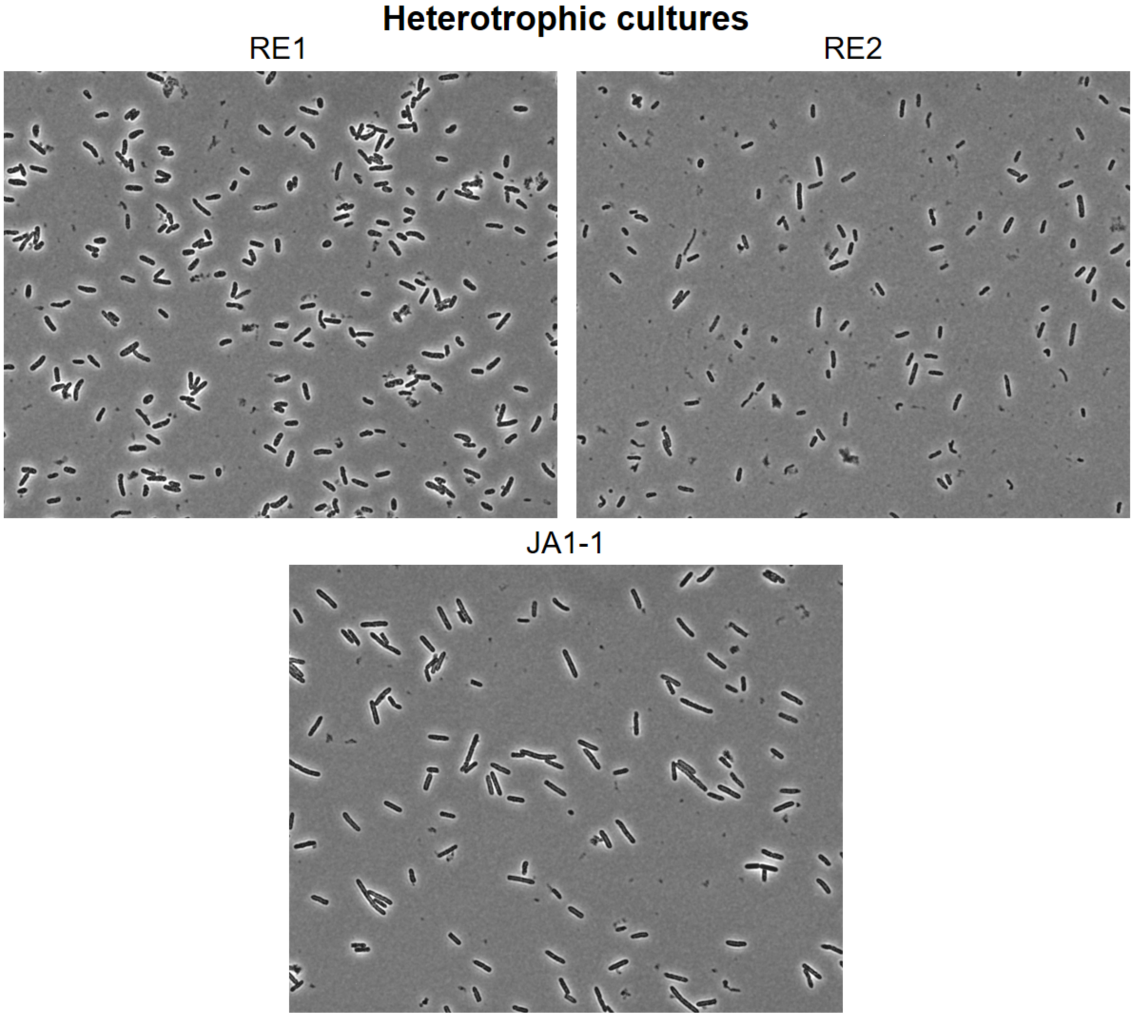


**Supplementary Fig. 15.** Micrographs showing lack of spores or sporulating cells in heterotrophic cultures of JA1-1, RE1, and RE2 after ~20 days of incubation. Micrographs taken regularly during the incubation period showed same results. Spores or endospores are expected to be white.

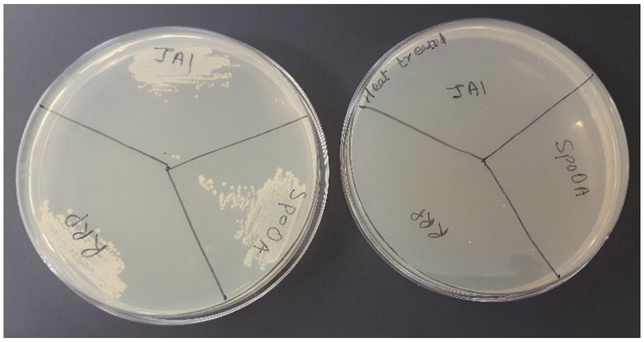


**Supplementary Fig. 16.** Lack of heat-resistant spores in heterotrophic cultures of JA1-1, RE1, and RE2. Left plate, cultures plated before heat-shock; right plate, cultures plates after heat-shock. JA1, JA1-1; Spo0A, RE1; RRP, RE2.

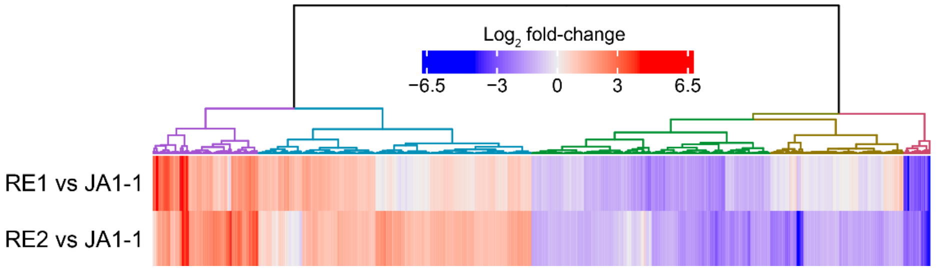


**Supplementary Fig. 17.** Hierarchical clustering of expression changes for all 583 differentially expressed proteins.
